# Genetic targeting of *Card19* is linked to disrupted *Ninj1 expression*, impaired cell lysis, and increased susceptibility to *Yersinia* infection

**DOI:** 10.1101/2021.03.19.436207

**Authors:** Elisabet Bjanes, Reyna Garcia Sillas, Rina Matsuda, Benjamin Demarco, Timothée Fettrelet, Alexandra A. DeLaney, Opher S. Kornfeld, Bettina L. Lee, Eric M. Rodriguez Lopez, Daniel Grubaugh, Meghan A. Wynosky-Dolfi, Naomi H. Philip, Elise Krespan, Dorothy Tovar, Leonel Joannas, Daniel P. Beiting, Jorge Henao-Mejia, Brian C. Schaefer, Kaiwen W. Chen, Petr Broz, Igor E. Brodsky

## Abstract

Cell death plays a critical role in inflammatory responses. During pyroptosis, inflammatory caspases cleave Gasdermin D (GSDMD) to release an N-terminal fragment that generates plasma membrane pores that mediate cell lysis and IL-1 cytokine release. Terminal cell lysis and IL-1β release following caspase activation can be uncoupled in certain cell types or in response to particular stimuli, a state termed hyperactivation. However, the factors and mechanisms that regulate terminal cell lysis downstream of GSDMD cleavage remain poorly understood. In the course of studies to define regulation of pyroptosis during *Yersinia* infection, we identified a line of *Card19*-deficient mice (*Card19^lxcn^)* whose macrophages were protected from cell lysis and showed reduced apoptosis and pyroptosis, yet had wild-type levels of caspase activation, IL-1 secretion, and GSDMD cleavage. Unexpectedly, CARD19, a mitochondrial CARD-containing protein, was not directly responsible for this, as two independently-generated CRISPR/Cas9 *Card19* knockout mice showed no defect in macrophage cell lysis, and expression of CARD19 in *Card19^lxcn^* macrophages did not restore cell lysis. *Card19* is located on chromosome 13, adjacent to *Ninj1*, which was recently reported to regulate cell lysis downstream of GSDMD activation. Intriguingly, RNA-seq and western blotting revealed that *Card19^lxcn^* BMDMs are hypomorphic for NINJ1 expression, and reconstitution of *Ninj1* in *Card19^lxcn^* immortalized BMDMs restored cell lysis. *Card19^lxcn^* mice exhibited significantly increased susceptibility to *Yersinia* infection, demonstrating that cell lysis itself plays a key role in protection against bacterial infection. Our findings identify genetic targeting of *Card19* being responsible for off-target effects on the adjacent *Ninj1* gene, thereby disrupting the ability of macrophages to undergo plasma membrane rupture downstream of gasdermin cleavage and impacting host survival and bacterial control during *Yersinia* infection.

**Author Summary:** Programmed cell death is critical for regulating tissue homeostasis and host defense against infection. Pyroptosis is an inflammatory form of programmed cell death that couples cell lysis with release of inflammatory cytokines. Cell lysis is triggered by activation of particular intracellular pore forming proteins, but how regulation of cell lysis occurs is not well understood. Genetic targeting of *Card19* on chromosome 13 resulted in decreased expression of the adjacent gene, *Ninj1* which was recently found to regulate terminal lysis events in response to cell death-inducing stimuli. We found that macrophages from *Card19*-deficient mice were resistant to multiple forms of cell death in response to a variety of inflammatory stimuli, including canonical and non-canonical inflammasome activation, as well as triggers of cell-extrinsic apoptosis. Notably, *Card19*-deficient mice were more susceptible to *Yersinia* infection, indicating that cell lysis contributes to control of bacterial infections. Our data provide new insight into the impact of terminal cell lysis on control of bacterial infection and highlight the role of additional factors that regulate lytic cell death downstream of gasdermin cleavage.

## Introduction

Regulated cell death is an evolutionarily conserved mechanism by which multicellular organisms control fundamental biological processes ranging from tissue development to microbial infection. Apoptosis and pyroptosis represent two distinct forms of regulated cell death that are activated in response to diverse stimuli, including developmental signals (1, 2), tissue stress or injury (3), and microbial infection (4–7). While apoptosis is an immunologically quiescent or suppressive form of cell death during tissue homeostasis and development, certain stimuli, including infections and chemotherapeutic agents, induce apoptosis that is accompanied by inflammatory signals that contribute to anti-pathogen and anti-tumor immunity (8–10). Apoptosis is activated by specific apoptotic initiator caspases, following certain intrinsic or extrinsic signals, and mediates organized disassembly of the cell via a process that limits cellular permeability and enables phagocytosis of the dying cell (11, 12). Alternatively, pyroptosis is a lytic form of regulated cell death characterized by caspase-1- or −11-dependent plasma membrane disruption mediated by cleavage and activation of the pore forming protein Gasdermin D (GSDMD), and release of interleukin-1 (IL-1) family cytokines and other intracellular components (13–16). However, inhibition of the receptor-proximal signaling kinases TAK1 or IKK by chemotherapeutic drugs or during infection by bacterial pathogens such as *Yersinia*, can trigger a caspase-8-dependent apoptosis pathway that is also associated with GSDMD cleavage (17–19). Interestingly, a recent study demonstrated that only caspase-8 activated in the RIPK1-containing complex IIb can efficiently cleave GSDMD in macrophages (20). These findings suggest a potential point of intersection of cell death pathways previously viewed as distinct (21).

During pyroptosis, secretion of IL-1 family cytokines and release of intracellular contents are temporally and genetically linked, and it has been suggested that IL-1 cytokines are released from cells when they undergo caspase-1 or −11-dependent lysis (22, 23). However, IL-1 secretion and cell lysis can be uncoupled in certain cell types or in response to specific stimuli. For example, the osmoprotectant glycine prevents release of lactate dehydrogenase (LDH), a key indicator of cell lysis, but does not affect IL-1β cytokine release (24, 25). Conversely, genetic ablation of IL-1 does not prevent cell lysis following pyroptotic stimuli. Moreover, IL-1 secretion can occur by living cells in the absence of apparent cytotoxicity (26, 27). Notably, neutrophils and monocytes display evidence of caspase-1 processing and IL-1 secretion in the absence of cell lysis (28, 29). Furthermore, dendritic cells treated with the oxidized lipid oxPAPC, or macrophages treated with the *N-*acetyl glucosamine fragment of bacterial peptidoglycan, release IL-1 without undergoing cell death (30, 31). Interestingly, GSDMD is genetically required for the release of IL-1 cytokines in response to non-pyroptotic stimuli, a state termed hyperactivation (26, 31, 32). Collectively, these data imply that while GSDMD processing and membrane insertion are critical for ultimate cell lysis, a cell fate decision checkpoint exists that distinguishes GSDMD-dependent IL-1 secretion from terminal cell death.

In our efforts to identify regulators of cell death during bacterial infection, we investigated the caspase activation and recruitment domain (CARD)-containing protein CARD19 (33). CARD19 is a mitochondrial membrane protein that contains an N-terminal CARD and C-terminal transmembrane domain, suggesting that it could be involved in the regulation of cell death or inflammatory responses (33, 34). Multiple mitochondria-associated proteins regulate inflammasome activation and inflammatory signaling (35–37). Notably, the mitochondrial CARD-containing protein, MAVS, plays a critical role in anti-viral immune signaling and IL-1 cytokine release (35, 38), and the mitochondrial outer membrane lipid cardiolipin plays an important role in regulating NLRP3 inflammasome assembly (39).

Intriguingly, primary macrophages from *Card19*-deficient mice (*Card19^lxcn^*) were resistant to cell lysis and release of lytic cell markers in response to apoptotic and pyroptotic stimuli, but exhibited wild-type levels of caspase activation, IL-1 secretion, and gasdermin processing. Unexpectedly however, two independent CRISPR/Cas9 knockout mouse lines that we generated did not show the same phenotype; furthermore, expression of CARD19 in immortalized BMDMs from the original *Card19^lxcn^* line did not restore their ability to undergo lysis, indicating that CARD19 itself is unlikely to be responsible for regulating terminal cell lysis in response apoptotic and pyroptotic stimuli. The cell lysis defect in *Card19^lxcn^* mice was unrelated to Sterile alpha and heat armadillo motif-containing protein (SARM1), which was reported to regulate NLRP3 inflammasome-dependent cell death independently of IL-1β cytokine release (40), as BMDMs from four independent *Sarm1*^-/-^ lines showed no defects in cell death, IL-1β release, and GSDMD cleavage. RNA-seq identified a handful of differentially regulated genes in *Card19^lxcn^* BMDMs, including a recently reported regulator of plasma membrane lysis, *Ninj1* (41), which is located adjacent to *Card19* on chromosome 13. *Card19^lxcn^* BMDMs showed significantly reduced *Ninj1* mRNA expression at baseline and substantially diminished NINJ1 protein levels both at baseline and upon LPS stimulation. Critically, reconstitution of immortalized *Card19^lxcn^* BMDMs rescued the defect in cell lysis, similar to *Ninj1^-/-^* BMDMs, indicating that the phenotype of *Card19^lxcn^* mice results from a defect in NINJ1 protein expression. *Card19^lxcn^* mice were more susceptible to infection with *Yersinia*, exhibiting enhanced mortality and systemic bacterial burdens relative to wildtype mice, implicating NINJ1-mediated lysis in host-pathogen interactions. Our data extend recent findings describing NINJ1 as a regulator of cell lysis downstream of caspase activation and gasdermin cleavage, and highlight the role of lytic cell death in host defense against bacterial infection.

## Results

### *Card19^lxcn^* BMDMs are deficient for caspase-dependent cell death

Our efforts to understand how cell death is regulated during bacterial infection led us to investigate the mitochondrial CARD-containing protein, CARD19, formerly known as BinCARD-2 (33, 34). To test the possible contribution of CARD19 to cell death during pyroptosis, we compared the kinetics of cell death in primary bone marrow derived macrophages (BMDMs) from B6 mice and mice with a targeted deletion in *Card19* generated by Lexicon Genetics (designated *Card19^lxcn^*), in response to infection by *Salmonella enterica* serovar Typhimurium (*S*. Tm) or LPS+ATP treatment. *S*. Tm infection of BMDMs activates the well-characterized NAIP/NLRC4/caspase-1 inflammasome pathway, leading to pyroptosis and IL-1 cytokine release (4, 5, 42, 43). In contrast, LPS+ATP treatment activates the NLRP3/ASC/caspase-1 inflammasome, which induces pyroptosis and IL-1 cytokine release via activation of the P2X7 receptor and potassium efflux (44–48). *Card19^lxcn^* BMDMs displayed a striking and significant defect in cell death following *S.* Tm infection or LPS+ATP treatment, as measured by release of LDH **(Fig. 1A and 1B)**. Consistently, *Card19^lxcn^* BMDMs exhibited significantly reduced levels of membrane permeability as determined by PI uptake, both in kinetics and amplitude **(Fig. 1C and 1D)**.

**Fig. 1.**
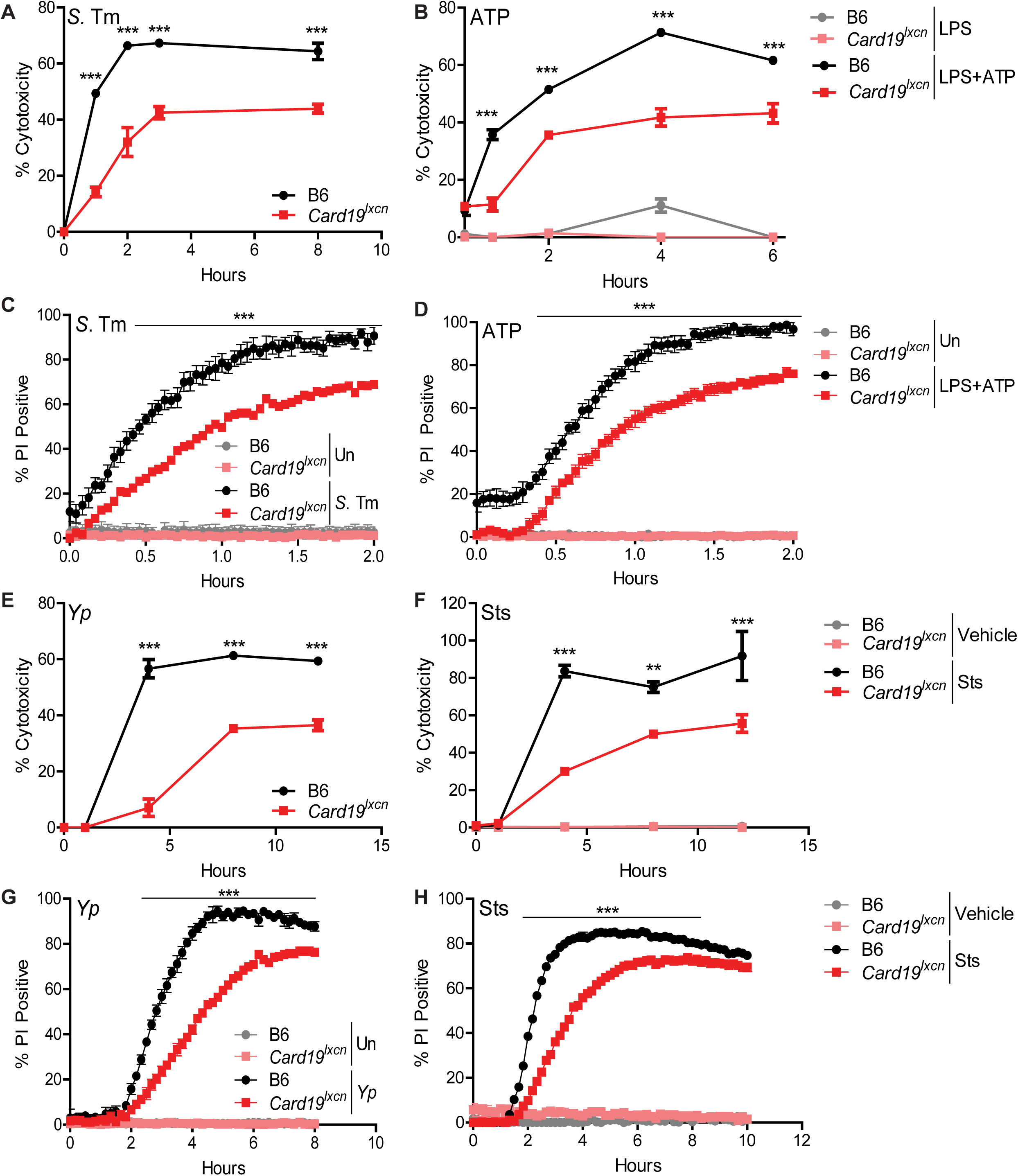
*Card19^lxcn^* BMDMs are deficient for caspase-dependent cell death. Primary C57BL/6J (B6) or *Card19^lxcn^* BMDMs were infected with *Salmonella* Typhimurium (*S*. Tm) or *Y. pseudotuberculosis* (*Yp*), or treated with LPS+ATP or staurosporine (Sts), as indicated, and the kinetics of cell death was assayed by release of lactate dehydrogenase (LDH) (A, B, E, F) or propidium iodide (PI) uptake (C, D, G, H). Each figure is representative of three or more independent experiments. LDH release was assayed at specific times post-infection. PI uptake was measured over a two or ten-hour timecourse with fluorometric measurements taken at 2.5 minute (C, D) or 10 minute (G, H) intervals. Mean ± SEM of triplicate wells is displayed. Each panel is representative of three or more independent experiments. *** p < 0.001, ** p < 0.01, * p < 0.05. n.s. not significant, analyzed by 2-way ANOVA with Bonferroni multiple comparisons post-test.

*Card19^lxcn^* BMDMs also exhibited a significant defect in cell death following infection by *Yersinia pseudotuberculosis* (*Yp*), which activates cell-extrinsic caspase-8- and RIPK1-dependent apoptosis in response to *Yp* blockade of IKK- and MAPK-signaling (6, 49, 50), as well as in response to staurosporine (Sts), a broad-spectrum kinase inhibitor that induces cell-intrinsic apoptosis and also activates caspase-8 in myeloid cells (51–53) **(Fig. 1E and 1F**). These observations were also mirrored by delayed PI uptake in *Card19^lxcn^* BMDMs **(Fig. 1G and 1H)**. Strikingly, *Card19^lxcn^* BMDMs were resistant to cell death in response to multiple apoptotic and pyroptotic stimuli, including cycloheximide, etoposide, and noncanonical inflammasome activation **(Fig. 2A-2C)**. However, RIPK3-dependent programmed necrosis which occurs in response to LPS and the pan-caspase inhibitor z-VAD-FMK (54, 55), was unaffected by CARD19-deficiency **(Fig. 2D)**. Altogether, these findings demonstrate that BMDMs from *Card19^lxcn^* mice exhibit increased resistance to membrane permeability and terminal cell lysis downstream of apoptotic and pyroptotic triggers.

**Fig. 2.**
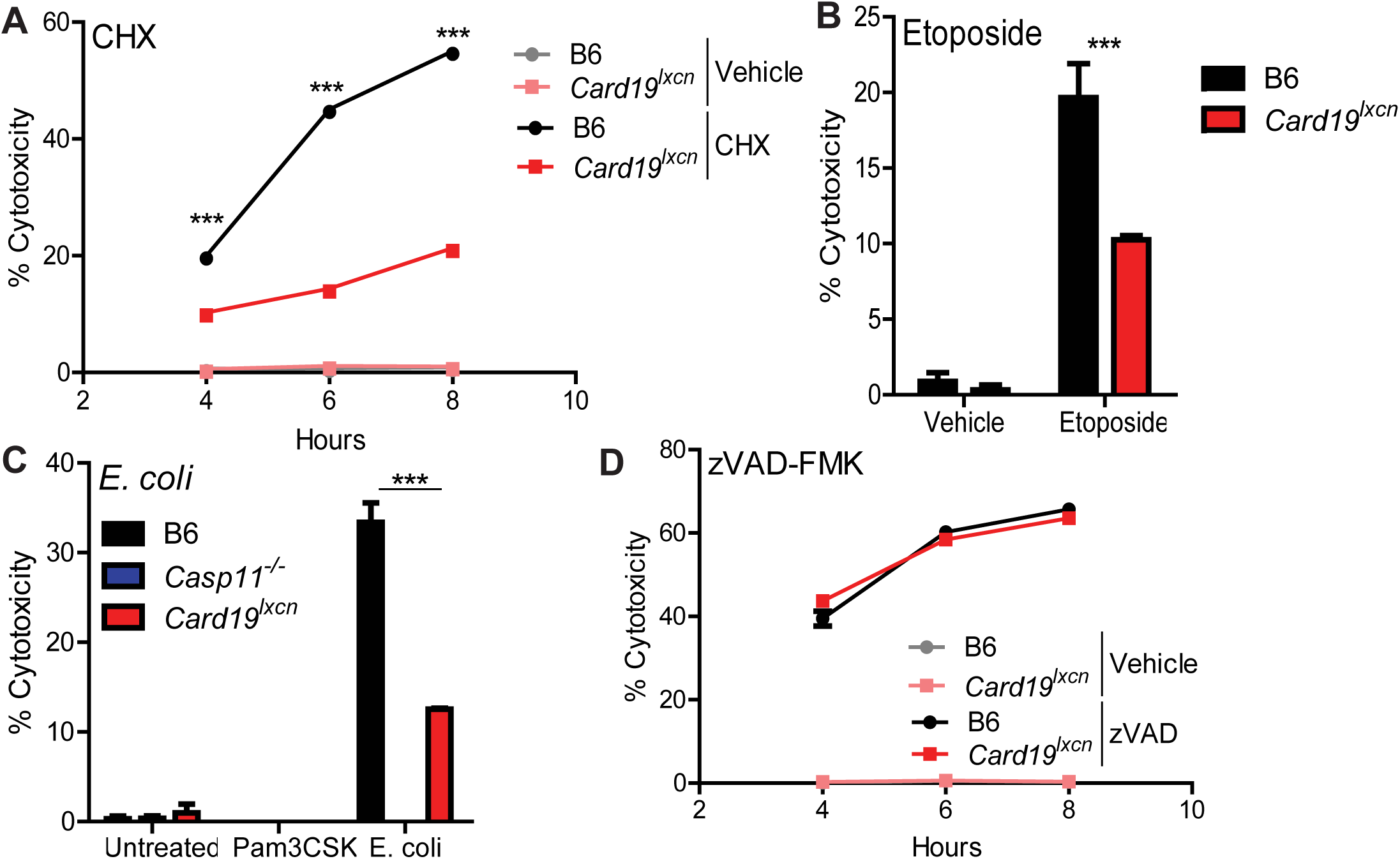
*Card19^lxcn^* BMDMs are caspase-dependent cell death but not necroptosis. Primary C57BL/6J (B6), *Card19*^lxcn^, *Card19^+/+^*, and *Casp11^-/-^* BMDMs were treated with (A) Cycloheximide (CHX) (B) etoposide, (C) Pam3CSK+*E. coli* or (D) z-VAD-FMK + LPS and cell death was assayed by LDH release. BMDMs were treated, supernatants were harvested from triplicate wells at indicated time points and measured for cytotoxicity. Mean ± SEM of triplicate wells is displayed. Each figure is representative of 2 or more independent experiments. *** p < 0.001, ** p < 0.01, * p < 0.05. n.s. not significant. 2-way ANOVA with Bonferroni multiple comparisons post-test.

### Peritoneal macrophages from *Card19^lxcn^* mice have reduced levels of cell death

The peritoneal cavity contains two distinct populations of macrophages, termed Large and Small (LPM and SPM, respectively)(56). At baseline, the primary population is the LPM, while 2 days post-injection of LPS or other inflammatory triggers, such as thioglycolate, the primary population in the peritoneal cavity shifts to the SPM (56, 57). Interestingly, thioglycolate-elicited peritoneal macrophages exhibited significantly higher levels of LDH than peritoneal macrophages isolated from PBS-treated mice following infection with either *S.*Tm or *Yp* **(Fig. 3A-3C)**. These data indicate that SPM undergo elevated levels of cell death relative to LPMs. Moreover, thioglycolate-elicited PMs from *Card19^lxcn^* mice had significantly lower levels of LDH release that C57BL/6J PMs, indicating that, similarly to BMDMs, *ex vivo* isolated macrophages from *Card19^lxcn^* animals also have a defect in cell lysis **(Fig. 3D and 3E)**.

**Fig. 3.**
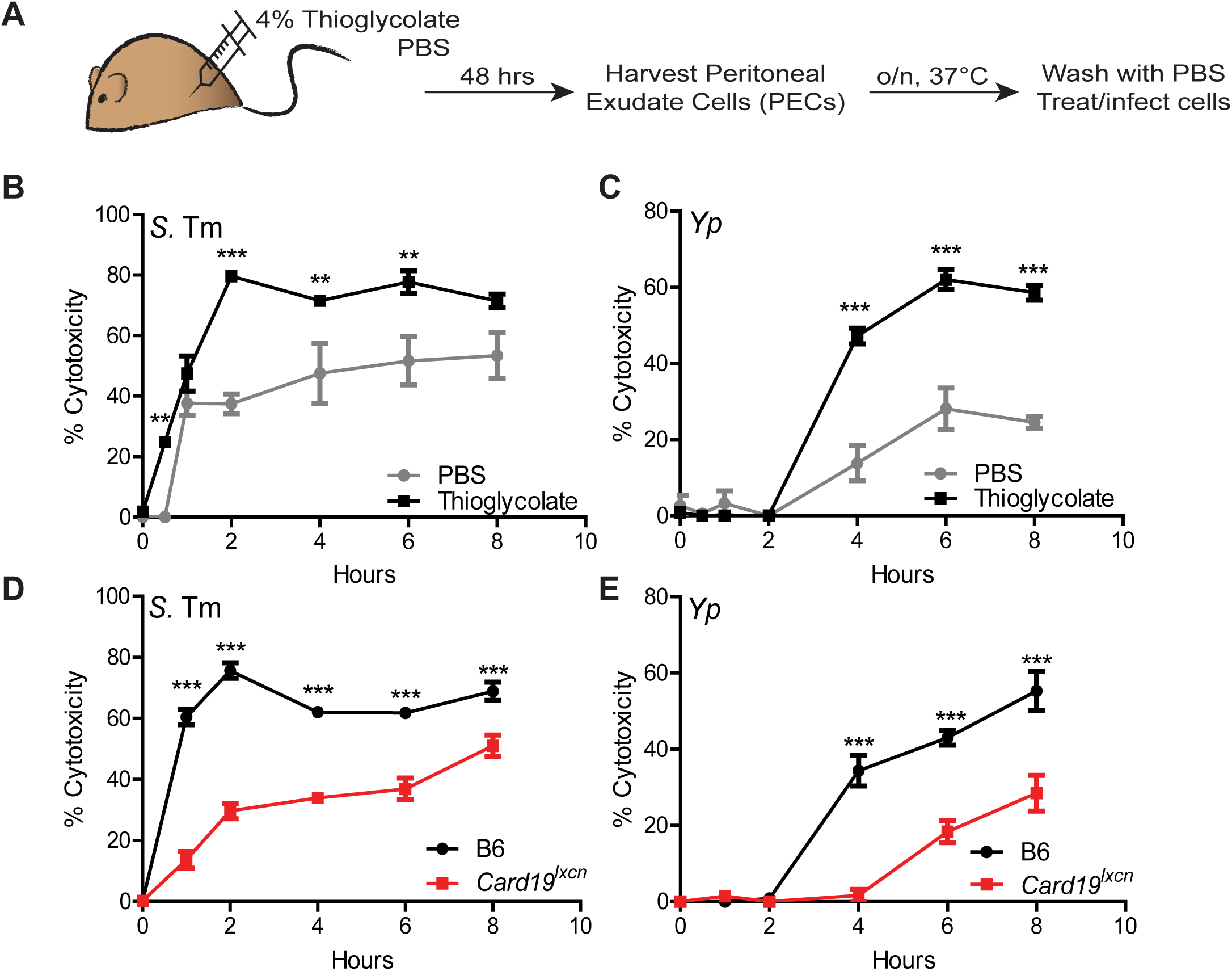
Peritoneal macrophages from *Card19^lxcn^* mice have reduced levels of cell death. (A) B6 and *Card19^lxcn^* mice were injected with 4% aged thioglycolate or PBS. 48 hrs later, PECS were harvested, RBC lysed, counted and plated in triplicate overnight at 37°C. PBS PECS were pooled prior to plating. Cells were washed with PBS to remove non-adherent cells and infected with *Yp* or *S.* Tm (MOI 10) or treated with staurosporine (10 uM). Cytotoxicity was measured by LDH release at indicated time points. *** p < 0.001, ** p < 0.01, * p < 0.05. n.s. not significant. 2-way ANOVA with Bonferroni multiple comparisons post-test. Graphs are representative of two independent experiments with 3-6 mice per group, per genotype. (B, C) PBS: mean ± SEM for technical replicates of 3 pooled B6 mice. Thioglycolate: means of technical replicates for 3 B6 mice ± SEM. Cells were infected with (B) *S.* Tm or (C) *Yp*. (D, E) PBS: mean ± SEM for technical replicates of 4-6 pooled B6 and *Card19*^lxcn^ mice + SEM. Thioglycolate: means of technical replicates for 4 B6 and *Card19^lxcn^* mice ± SEM. Cells were infected with (B) *S.* Tm or (C) *Yp*.

### *Card19^lxcn^* BMDMs retain intracellular alarmin HMGB1 following activation of cell death

HMGB1 is a nuclear chromatin-associated protein that is released from cells upon loss of plasma membrane integrity (58). As expected in B6 cells, *S*. Tm or LPS+ATP treatment led to significant HMGB1 release into the supernatant by one-hour post-infection, and by 4 hours post-exposure to *Yp* or staurosporine (Sts) **(Fig. 4A)**. In contrast, *Card19^lxcn^* cells exhibited significantly delayed release of HMGB1 for each respective timepoint, consistent with our findings that they exhibit increased resistance to membrane rupture **(Fig. 4A)**. HMGB1 was virtually undetectable in *Card19^lxcn^*macrophage supernatants one-hour post-treatment with *S*. Tm or LPS+ATP, and minimally detectable at 4 hours post-treatment in response to *Yp* or Sts **(Fig. 4A)**. To obtain insight into the dynamics of HMGB1 release from cells, we analyzed HMGB1 release by confocal microscopy following exposure of Wt and *Card19^lxcn^* cells to Sts. We observed three morphologies of HMGB1 staining – nuclear, in which the majority of HMGB1 was located in the nucleus; cellular, in which HMGB1 was absent from the nucleus but still contained within the confines of a cell; and extracellular, in which a cloud of HMGB1 was detectable surrounding what was likely the remnants of apoptotic cells and cellular debris. Notably, and consistent with intracellular retention of HMGB1 in *Card19^lxcn^* cells, a higher frequency of *Card19*^lxcn^ cells relative to B6 BMDMs retained nuclear HMGB1 **(Fig. 4B and 4C)**, while B6 BMDMs exhibited extracellular HMGB1 cloud formation much more frequently than *Card19^lxcn^* BMDMs **(Fig. 4B white arrows and quantified in Fig. 4D)**. Altogether, these findings indicate that *Card19^lxcn^* BMDMs resist cell lysis and release of intracellular contents during induction of cell death triggered by caspase-1 or caspase-8.

**Fig. 4.**
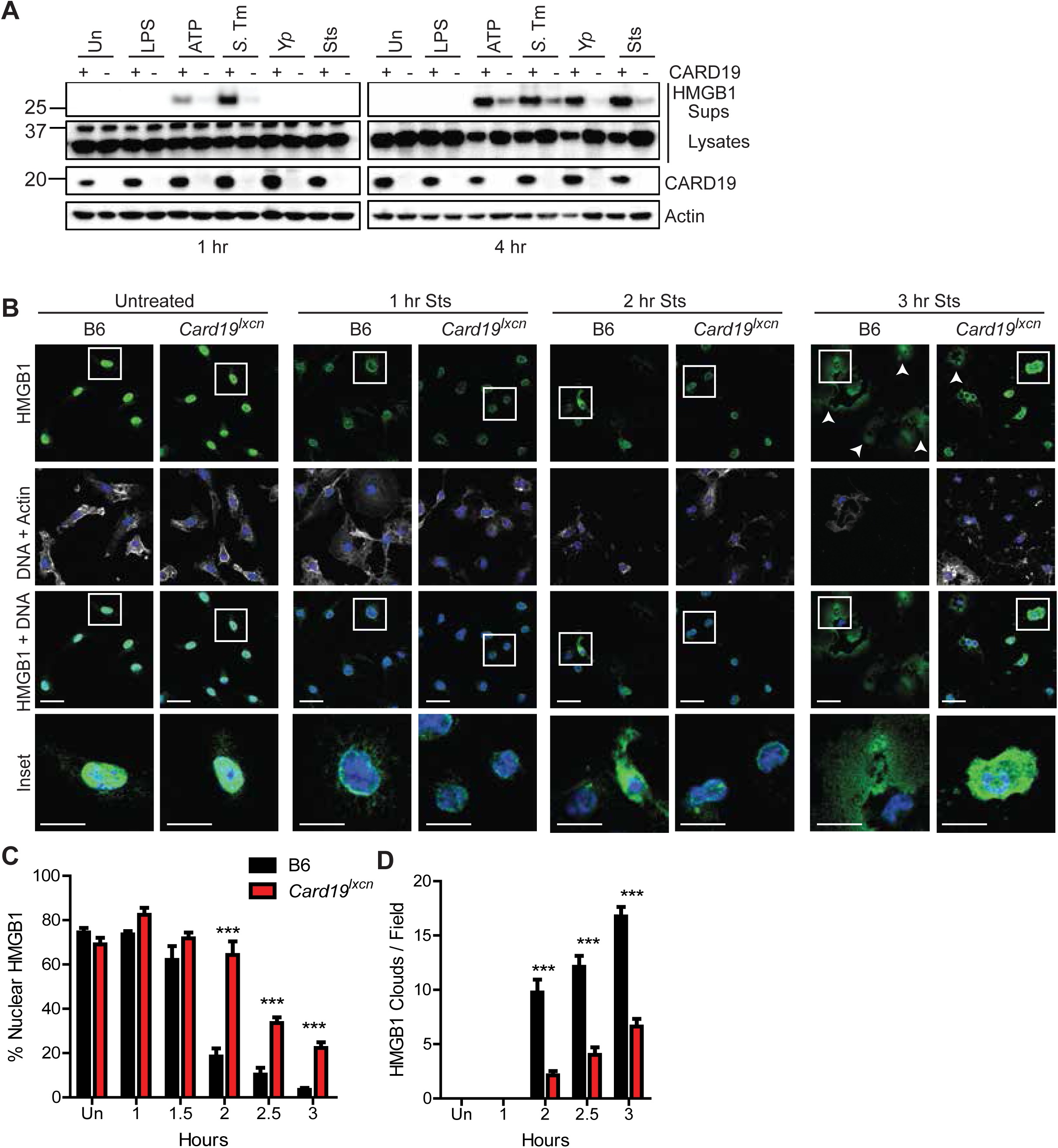
*Card19^lxcn^* BMDMs retain intracellular alarmin HMGB1 following activation of cell death. (A) Primary BMDMs from B6 (+) or *Card19^lxcn^* (-) mice were treated with indicated treatments or infections, and TCA precipitated supernatants (Sups) or whole cell lysates (Lysates) were harvested at 1 or 4 hours post-infection and analyzed by western blotting for HMGB1, CARD19, and actin (loading control). ATP indicates cells that were primed with LPS for 3 hours and treated with ATP for 1 or 4 hours. *S.Tm – Salmonella* Typhimurium; *Yp Yersinia pseudotuberculosis.* Sts – staurosporine. (B) Confocal microscopy images of untreated and staurosporine-(Sts) treated B6 and *Card19*^lxcn^ BMDMs fixed at indicated times post-Sts treatment and stained for HMGB1, Actin and DNA (Hoescht). White arrows indicate HMGB1 clouds. Scale bar 20 microns, inset 10 microns. (C) Quantification of nuclear HMGB1. n=25-50 cells per field, 5-8 fields per condition, per timepoint. (D) Quantification of HMGB1 clouds in B6 and *Card19*^lxcn^ BMDMs after staurosporine treatment. 5-8 fields quantified per condition, per timepoint. Mean ± SEM is displayed. *** p < 0.001, ** p < 0.01, * p < 0.05. n.s. not significant. 2-way ANOVA with Bonferroni multiple comparisons post-test. Representative of 3 (A-C) or 2 (D) independent experiments.

### *Card19^lxcn^* macrophages are not deficient in activation of caspase-1 or −8

Most currently known regulators of cell death act at the level of caspase activation or assembly of caspase-activating inflammasome complexes (59). Pyroptosis is mediated by activation of caspase-1 or −11 (15, 59, 60), which cleave Gasdermin D (GSDMD), thereby triggering cell lysis by enabling formation of oligomeric N-terminal GSDMD pores in the plasma membrane (14, 16, 61, 62). Surprisingly, despite the significant reduction in cytotoxicity in *Card19^lxcn^* cells, processing of caspase-1 was equivalent in B6 and *Card19^lxcn^* cells 15 minutes post-infection, at which time cell death is virtually undetectable in either genotype **(Fig. 5A)**. Moreover, B6 cells released significant amounts of cleaved and pro-caspase-1 into cell supernatants at later timepoints, in contrast to *Card19^lxcn^* BMDMs **(Fig. 5A)**, consistent with the reduced levels of pyroptosis in *Card19^lxcn^* cells in response to *S*. Tm infection. Reduced levels of cleaved caspase-1 in *Card19^lxcn^* BMDM supernatants was matched by increased amounts of cleaved caspase-1 in *Card19^lxcn^* whole cell lysates **(Fig. 5A)**. Similarly, initial processing of GSDMD to the cleaved p30 form was equivalent in B6 and *Card19^lxcn^* cell lysates, while the release of processed GSDMD into the supernatant was significantly reduced in *Card19*^lxcn^ cells **(Fig. 5A)**. Thus, *Card19^lxcn^* cells have no defect in either caspase-1 activation, or in the cleavage of caspase-1 targets.

**Fig. 5.**
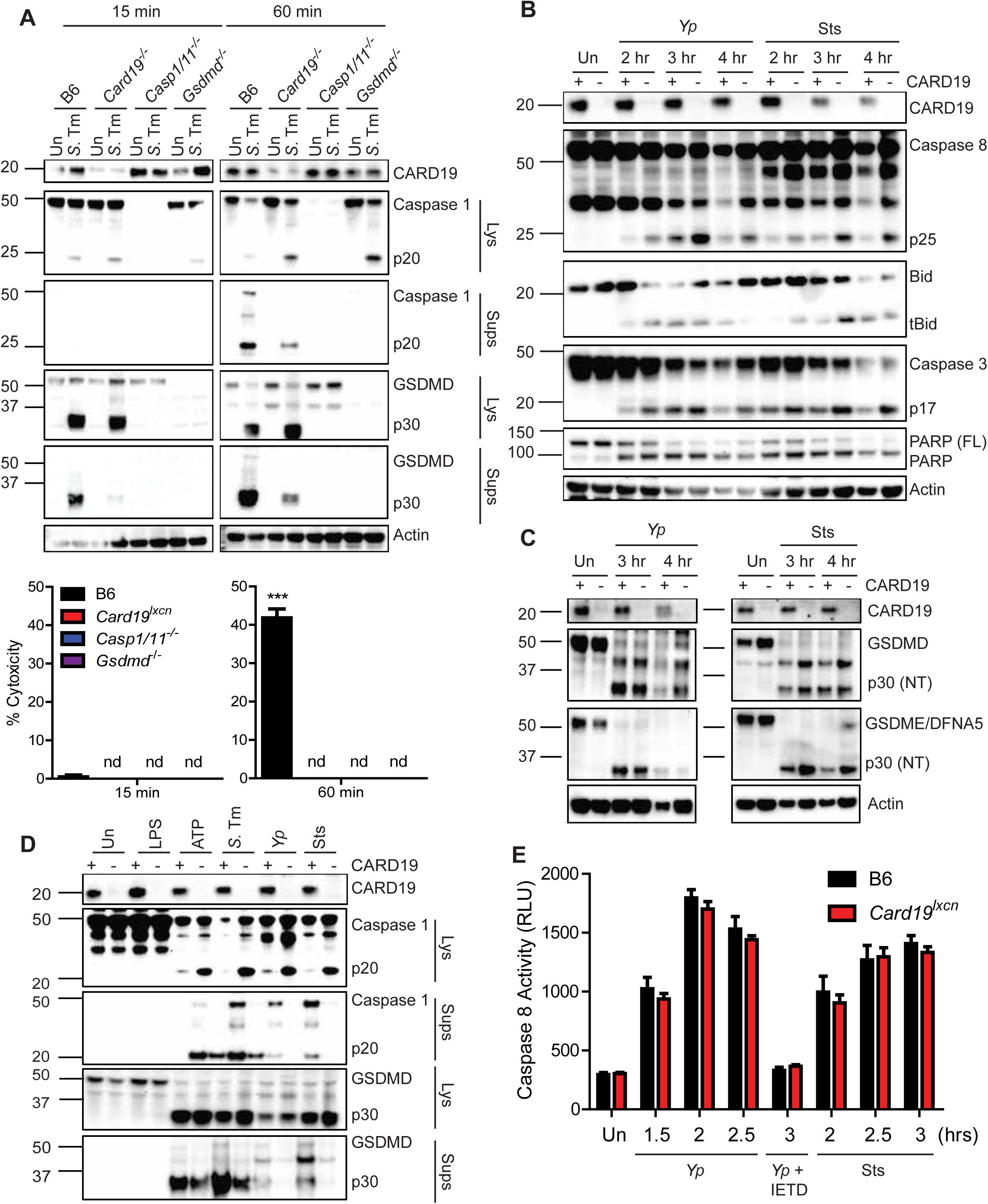
*Card19^lxcn^* macrophages are not deficient in activation of caspase-1 or −8. (A) B6, *Card19*^lxcn^, *Casp1*/*11*^-/-^ or *Gsdmd*^-/-^ BMDMs were left untreated or infected with *S*. Tm, and cell lysates (Lys) and TCA-precipitated supernatants (Sups) were run on SDS-PAGE and analyzed by western blotting for cleaved Casp1, GSDMD, or Actin (cell lysate loading control). Cell death from these cells was assayed in parallel by LDH release (indicated below blots). Mean ± SEM is displayed. 2-way ANOVA with Bonferroni multiple comparisons post-test. *** p < 0.001, ** p < 0.01, * p < 0.05. n.d. not detectable. (B) B6 (+) or *Card19^lxcn^* (-) BMDMs were infected with *Yp* or treated with Sts or 2, 3, or 4 hours, or left untreated (Un) as indicated. Cell lysates were prepared at indicated times and analyzed for presence of CARD19, cleaved caspase-8, tBid, cleaved caspase-3, cleaved PARP, and Actin (loading control). (C) B6 and *Card19^lxcn^* BMDMs were left untreated, infected with *Yp* or treated with staurosporine. Lysates were harvested at the indicated time points, run on SDS-PAGE and analyzed by western blotting for CARD19, GSDMD, GSDME/DFNA5, and actin (loading control). (D) B6 (+) and *Card19*^lxcn^ (-) BMDMs were left untreated, treated with LPS, LPS+ATP, Sts, or infected with *S.* Tm or *Yp*. Cell lysates (Lys) and supernatants (Sups) were harvested 1 hour post-ATP or S.Tm infection or 3 hours post Sts treatment and *Yp* infection and assayed for CARD19, Caspase-1, and presence of cleaved GSDMD. (E) B6 and *Card19^lxcn^* BMDMs were left untreated, infected with *Yp*, or treated with staurosporine. BMDMs were pretreated with the caspase-8 inhibitor IETD for one hour prior to infection. Caspase-8 activity was assessed by Caspase-8-Glo. Mean ± SEM is displayed. Blots, caspase-8 activity and LDH are representative of at least 3 independent experiments.

Cell-extrinsic apoptosis is mediated by oligomerization and auto-processing of caspase-8 (63, 64). Active caspase-8 directly cleaves its pro-apoptotic downstream targets, including the Bcl-2 family member Bid (65, 66), and the executioner caspases-3 and −7. Similarly to caspase-1, cleavage of caspase-8 as well as the processing of caspase-8 substrates, including Bid, caspase-3, the caspase-3 target poly-ADP ribose polymerase (PARP), and the pore-forming protein GSDME also known as DFNA5 (18, 67), were equivalent in B6 and *Card19^lxcn^* cells in response to *Yp* infection or staurosporine treatment **(Fig. 5B and 5C)**. Notably, we observed GSDMD processing in response to *Yp* infection as well as Sts treatment, in both B6 and *Card19^lxcn^* BMDMs **(Fig. 5C)**. Importantly, direct comparison of caspase-1 and GSDMD processing in response to multiple stimuli revealed that *Card19^lxcn^* cells had reduced release of cleaved caspase-1 and N-terminal GSDMD into the supernatant, and correspondingly increased retention in cell lysates, consistent with reduced terminal lysis of the cells in the absence of CARD19 **(Fig. 5D)**. *Card19^lxcn^* BMDMs had wild-type levels of caspase-8 activity in response to either *Yp* infection or Sts treatment, confirming that *Card19^lxcn^* BMDM do not have a defect in caspase activity **(Fig. 5E)**. Altogether, these findings demonstrate that *Card19^lxcn^* BMDMs do not have a defect in caspase activation or downstream target cleavage, and therefore likely have a defect in cell lysis downstream of caspase activation.

How *Card19^lxcn^* cells might retain membrane integrity downstream of both inflammatory and apoptotic caspase activation is not clear. However, caspase-8 and caspase-1 can both cleave IL-1β (51, 68, 69), as well as GSDMD (17, 18, 20) raising the question of whether *Card19^lxcn^* cells and *Gsdmd^-/-^* cells have similar defects in cell lysis in response to *Yp* infection. Surprisingly, although *Gsdmd^-/-^* BMDMs exhibited a significant defect in cell death in response to *Yp* infection **(Fig. S1A)**, this defect was not as pronounced as in *Card19^lxcn^* BMDMs, suggesting that *Card19^lxcn^* cells likely have a defect in additional GSDMD-independent mechanisms of cell lysis, potentially involving GSDME/DFNA5, which was also processed in response to *Yp* infection **(Fig. S1B and S1C)**. Consistent with recent findings (17–19), we observed cleaved GSDMD in *Casp1/11^-/-^* lysates following either *Yp* infection or staurosporine treatment. Although levels of cleaved GSDMD were significantly lower than in Wt cells, this finding is consistent with findings that caspase-8 cleaves GSDMD less efficiently than caspase-1 **(Fig. S1B and S1C)** (**19**). The caspase-8-selective inhibitor, IETD, abrogated GSDMD and GSDME cleavage in response to *Yp* or staurosporine **(Fig. S1B and S1C)**, suggesting that *Yersinia* and staurosporine induce GSDMD and GSDME cleavage in *Casp1/11^-/-^* cells in a caspase-8-dependent manner. Consistently, *Ripk3^-/-^* BMDMs showed robust cleavage of GSDMD and GSMDE while *Ripk3^-/-^Casp8^-/-^* BMDMs showed a near-complete absence of N-terminal GSDMD and GSDME p30 following *Yp* infection and significant reduction in GSDMD and GSDME processing after staurosporine treatment **(Fig. S1B and S1C)**. Notably, caspase-1 activity was not defective in the *Ripk3^-/-^Casp8^-/-^* BMDMs **(Fig. S1D)**.

### The cell death defect in *Card19^lxcn^* BMDMs is independent from cytokine release

Consistent with our findings that *Card19^lxcn^* BMDMs are not defective in their ability to regulate proteolytic activity of caspase-1, CARD19-deficient cells released wild-type levels of IL-1 cytokines in response to *S*. Tm, LPS+ATP, or infection with the Δ*yopEJK Yp* strain, which triggers a caspase-11/NLRP3-dependent pathway of IL-1 cytokine release (70, 71) **(Fig. 6A, B)**. These data indicate that *Card19^lxcn^* cells most likely do not have a defect at the level of GSDMD-dependent pore formation and cytokine release and that the *Card19^lxcn^* cells decouples secretion of IL-1β from pyroptosis, which normally occurs in response to these stimuli (26, 32). Consistent with a recent finding that *Card19^lxcn^* cells do not have altered NF-κB activation (33), *Card19^lxcn^* BMDMs did not have a defect in secretion of IL-6, TNF, or IL-12p40, **(Fig. 6C-6E)**. Altogether, these findings suggest that *Card19^lxcn^* cells have decoupled cell lysis as determined by LDH and HMGB1 release and the release of caspase-1-dependent cytokines.

**Fig. 6.**
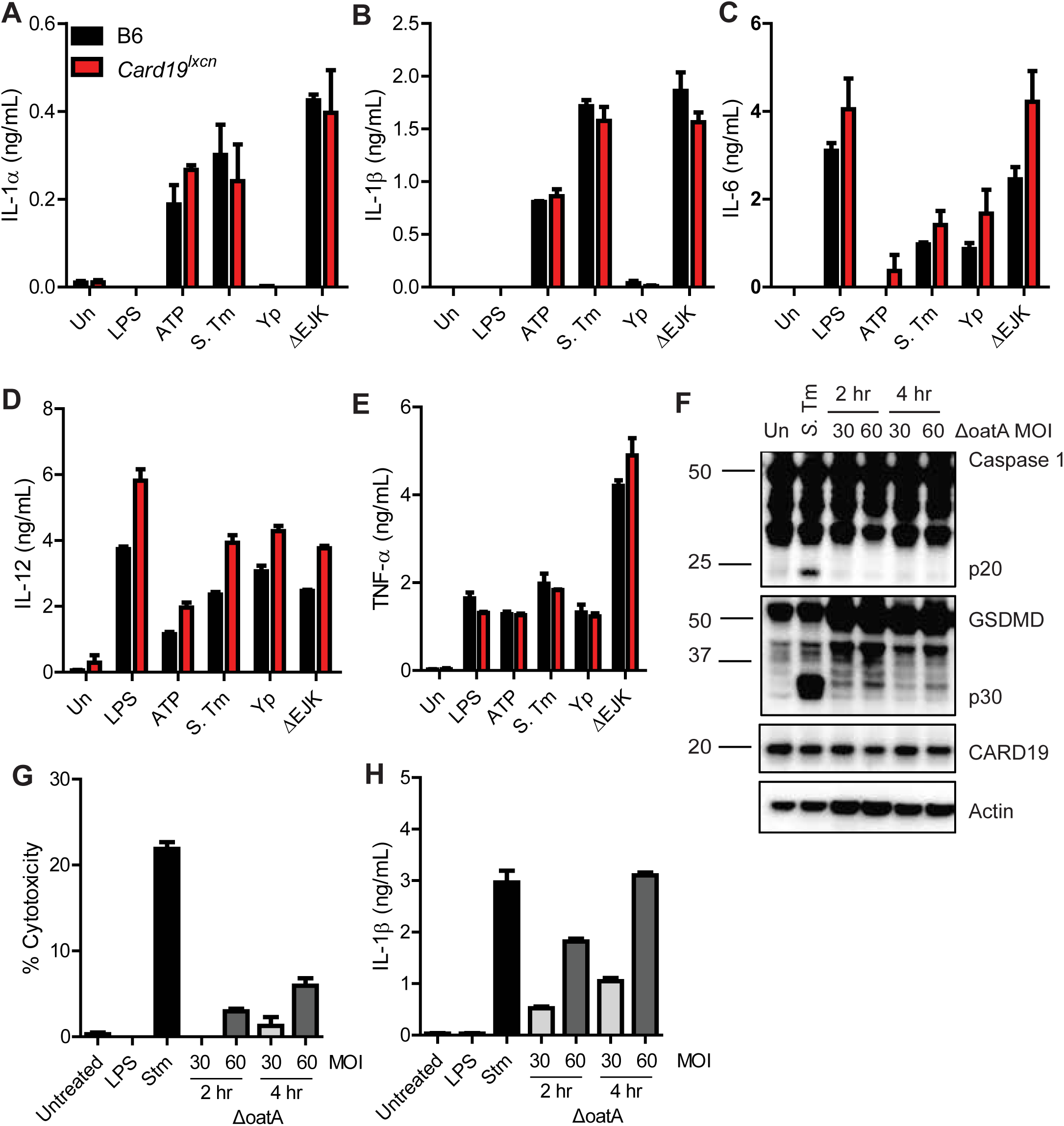
The cell death defect in *Card19^lxcn^* BMDMs is independent from cytokine release. (A-E) B6 (black bars) and *Card19*^lxcn^ (red bars) BMDMs were primed with LPS for 3 hours and treated with extracellular ATP, infected with *S*. Tm, *Yp*, or the *Yp* ΔEJK strain lacking effectors YopE, YopJ and YopK. Supernatants were harvested 2 hours post-infection or ATP treatment and analyzed by ELISA for release of (A) IL-1α, (B) IL-1B, (C) IL-6, (D) IL-12, and (E) TNF-α. Mean ± SEM is displayed. Each figure is representative of 3 or more independent experiments. (F-H) B6 BMDMs were primed with LPS for 4 hours and infected with Δ*oatA S. aureus* or *S.* Tm. (F) Lysates were harvested at indicated times, run on SDS-PAGE page and analyzed by western blotting for Caspase 1, GSDMD, CARD19, or Actin (loading control). Supernatants were harvested at indicated time points and analyzed by (G) LDH for cytotoxicity and (H) ELISA for release of IL-1β. Each figure is representative of two independent experiments.

Cells can release IL-1β while remaining viable during a state known as hyperactivation (26, 32, 72). GSDMD is required for IL-1β secretion during hyperactivation, but the mechanism by which GSDMD mediates this IL-1β release without inducing lysis is not understood (26, 72). *Staphylococcus aureus* lacking OatA induces hyperactivation (26, 72). Despite inducing robust IL-1β release under conditions of minimal LDH release, Δ*oatA S. aureus-*infected B6 BMDM exhibited minimal levels of caspase-1 and GSDMD processing **(Fig 6F-6H)**. CARD19 expression did not correlate with differences in hyperactivation exhibited between untreated, Δ*oatA* and *S.* Tm-infected BMDMs **(Fig. 6F)**. Since *Card19^lxcn^* BMDMs process caspase-1 and GSDMD normally while releasing IL-1 cytokines in the absence of overt lysis, these data indicate that lack of death in *Card19^lxcn^* cells is not evidence of a hyperactivation phenotype, but are likely resistant to cell lysis due to a defect at a terminal stage of pyroptosis. Curiously, Δ*oatA*-infected cells exhibited significantly lower levels of GSDMD processing, indicating that threshold levels of GSDMD processing or membrane insertion may underlie the distinction between hyperactivation and cell lysis **(Fig. 6F)**.

### *Card19^lxcn^* BMDMs are hypomorphic for *NINJ1* gene and protein expression

Numerous mitochondrial proteins have been implicated in recent years as essential regulators of caspase activity, GSDMD and terminal events in during lysis (35, 37, 73). While these studies were in progress, Sterile alpha and armadillo motif-containing protein (SARM1) was reported to regulate terminal cell lysis following stimulation of the NLRP3 inflammasome (40), and like CARD19, also localizes to the mitochondrial outer membrane (33). We therefore considered the possibility that SARM1 might contribute to cell lysis via interaction with CARD19. Unexpectedly however, BMDMs isolated from multiple independent lines of *Sarm1^-/-^* mice (described further in Materials and Methods), including *Sarm1^-/-^* mice reported by Carty *et al*. (40) and originally described by Kim et al (74), exhibited no defect in their ability to undergo cell lysis in response to canonical NLRP3 inflammasome stimuli **(Fig. S2A)**, nor did they exhibit elevated levels of inflammasome-dependent IL-1β release **(Fig. S2B).** *Sarm1^-/-^* BMDMs also showed wild-type levels of Caspase-1, GSDMD, and IL-1β processing **(Fig. S2C and S2D)**. Moreover, TNF release was unaffected in cells lacking SARM1 compared to WT controls, indicating that the NLRP3 inflammasome priming step (NF-κB pathway) is not dysregulated by the absence of SARM1 (**Fig. S2E**). Furthermore, in contrast to *Card19^lxcn^* BMDMs, the *Sarm1^-/-^* macrophage lines had wild-type levels of cell death and cytokine release in response to non-canonical inflammasome activation **(Fig. S2F and S2G)**. Notably, some *Sarm1^-/-^* lines were reported to contain a passenger mutation in the closely-linked gene *Xaf1*, indicating that some previous cell death-related phenotypes attributed to SARM1 may be due to alteration of *Xaf1* (75). Altogether, these findings indicate that the phenotype of *Card19^lxcn^* BMDMs is not related to potential interactions with SARM1.

Next, to test the sufficiency of CARD19 to induce cell death, we generated immortalized *Card19^+/+^* and *Card19^lxcn^* bone marrow hematopoietic progenitors using the ER-HOXB8 system (76), and transduced the progenitors with CARD19-expressing lentiviral constructs. Unexpectedly, *Card19^lxcn^* iBMDMs transduced with CARD19 displayed comparable death to vector control-transduced iBMDMs, raising questions as to whether CARD19-deficiency was directly responsible for the observed defect in caspase-dependent death in *Card19^lxcn^* BMDMs **(Fig. 7A)**.

**Fig. 7.**
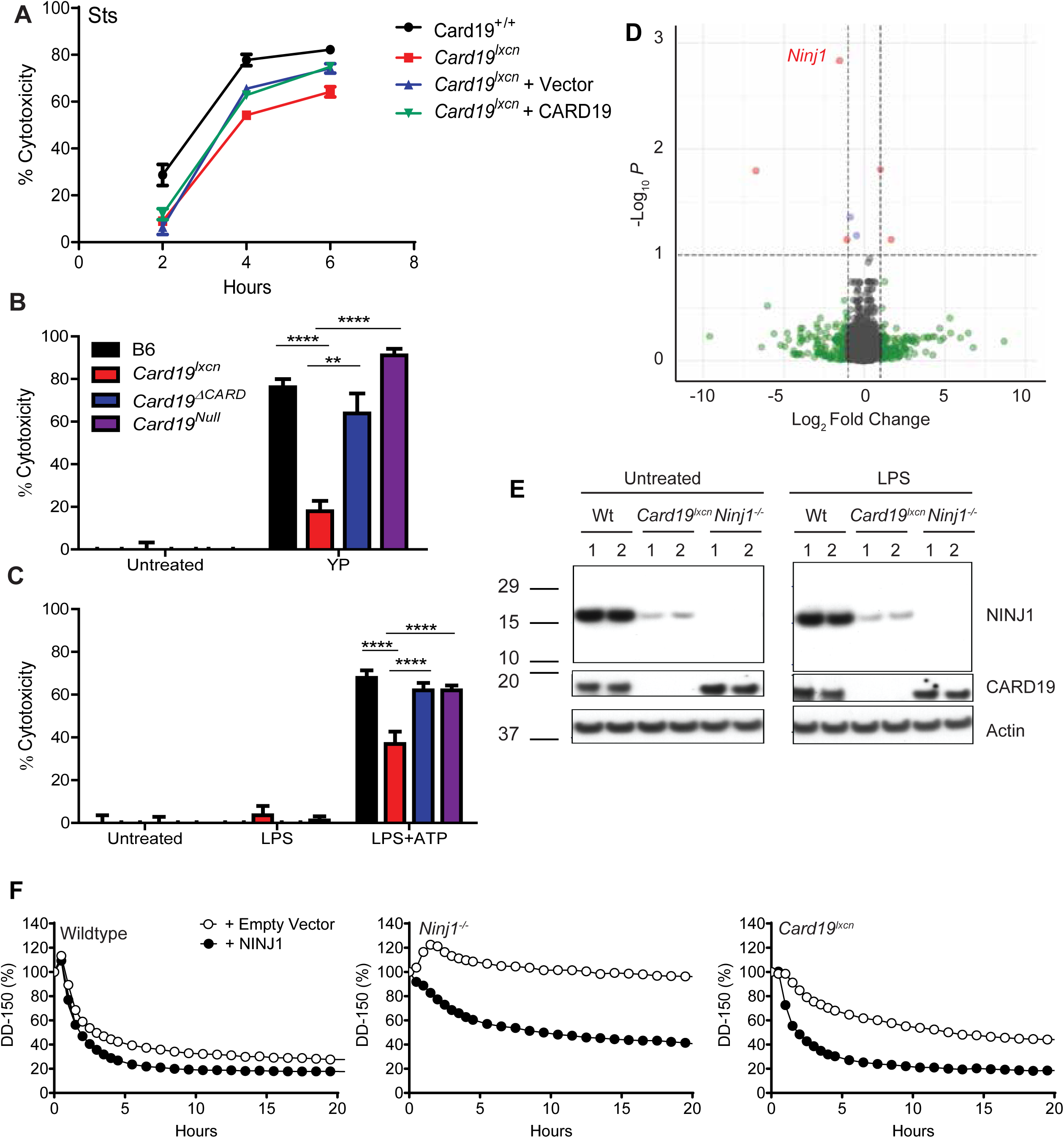
*Card19^lxcn^* BMDMs are hypomorphic for *Ninj1* and reconstitution with *Ninj1* restores cell death. (A) *Card19^lxcn^* immortalized murine progenitors were stably transduced with CARD19 or empty vector. Mature macrophages derived from untransduced immortalized progenitors *Card19^lxcn^* and Card19^+/+^ and transduced immortalized progenitors *Card19^lxcn^* + Vector and *Card19^lxcn^* + CARD19 were treated with sts. Cell death was assayed by LDH release at indicated time points. Representative of three independent experiments. (B,C) B6, *Card19^lxcn^*, *Card19^ΔCARD^*, and *Card19^Null^* BMDMs were treated with (B) *Yp* or (C) LPS+ATP. Cell death was assayed by LDH release. Representative of three independent experiments. 2-way ANOVA with Bonferroni multiple comparisons post-test. **** p < 0.0001 *** p < 0.001 ** p < 0.01, * p < 0.05, n.s. not significant. (D) Volcano plot from RNA-seq analysis on untreated B6 and *Card19^lxcn^* BMDMs showing differentially regulated genes. Genes whose expression is significantly altered are in red. *Ninj1* is marked. (E) BMDMs from B6, *Card19^lxcn^*, *Card19^ΔCARD^*, and *Card19^Null^* mice were primed with LPS or left untreated. Cell lysates were harvested, run on SDS-PAGE and analyzed by western blotting for cleaved CARD19, NINJ1, and Actin (cell lysate loading control). (F) Wildtype, *Ninj1^-/-^* and *Card19^lxcn^* iBMDMs were reconstituted with NINJ1 or empty vector. Release of dextran dye-150 (DD-150) in live cell imaging analysis following LPS electroporation over a 20 hour time course.

Passenger mutations in genes closely linked with the gene of interest in backcrossed C57BL/6 mice can confound interpretation of immune responses and cell death pathways in these lines (23, 77, 78). Notably, *Card19^lxcn^* mice were originally generated using 129SvEvBrd ESCs, and backcrossed for 10-12 generations (33). *Card19^+/lxcn^* and 129/SvIm/J BMDMs phenocopy wildtype BMDMs with respect to their ability to undergo terminal cell lysis in response to *S.* Tm., *Yp*, and Sts, indicating that the phenotype is linked to the *Card19* locus **(Fig. S3A-C)**. To rule the possibility that a passenger mutation linked to the *Card19* locus might be responsible, we next generated two independent *Card19*-deficient murine lines directly on the C57BL/6J background using CRISPR/Cas9. The first of these removed exons 2-3, removing the entire CARD domain, but leaving a truncated mRNA that could produce a peptide fragment containing the transmembrane domain, which we termed *Card19^ΔCARD^* **(Figure S3D, Table S1)**. The second was a complete deletion of the *Card19* locus, which we termed *Card19^Null^* (see Methods for further details of construction of these lines). Critically, *Card19^ΔCARD^* and *Card19^Null^* BMDMs exhibited no observable defect in their ability to undergo cell lysis following infection with either *Yp* or treatment with LPS+ATP, in contrast to BMDMs derived from the original *Card19^lxcn^* line **(Fig. 7B and 7C)**. Altogether, these data indicate that loss of CARD19 is not the mechanistic basis of the defect in cell death seen in the original *Card19^lxcn^* cells.

To identify the regions of the *Card19^lxcn^* chromosome that may contain the original 129SvEvBrd lineage, we performed SNP genotyping analysis of the *Card19^lxcn^* mice which indicated that approximately six megabases adjacent to the *Card19* locus were derived from the 129SvEvBrd lineage **(Fig. S4A, Table S2)**. We additionally performed whole exome sequencing on B6 and *Card19^lxcn^* mice to address the possibility that *Card19^lxcn^* mice contain a functionally important polymorphism or have an expression defect in a gene closely linked to Card19. Although whole exome sequencing identified potential mutations in eight genes on chromosome 13, they were not high probability candidates **(Table S3)**.

We further performed RNA-seq on B6 and *Card19^lxcn^* BMDMs, with the hypothesis that transcriptional alteration of genes closely linked to *Card19^lxcn^* might result in the observed loss of cell lysis **(Fig. 7D, Table S4)**. Intriguingly, *Ninj1*, recently identified as a key regulator of plasma membrane rupture during lytic cell death in response to apoptotic, pyroptotic and necrotic stimuli (41) is directly upstream of *Card19*, making it a likely candidate for passenger mutations or off-target effects **(Fig. S4A, blue arrow)**. *Card19^lxcn^* and *Ninj1^-/-^* BMDMs have a strikingly similar phenotype, as they both have a defect in cell lysis that is independent of IL-1 cytokine release and processing of caspases and gasdermins ((41), **Fig. 1-5**). SNP mapping with a high-density SNP array identified a 129SvEvBrd-derived region in *Card19^lxcn^* cells downstream of both *Ninj1* and *Card19* that also contained several B6 SNPs (**Table S2, Fig. S4A**). It is therefore likely that complex recombination occurred at the *Ninj1/Card19* locus. To test whether any mutations in *Ninj1* were present in the *Card19^lxcn^* mice, we directly sequenced all 5 exons of *Ninj1*, including the 3’ splice acceptor site for exon 2, reported to be mutated in the ENU-mutagenesis screen that first identified *Ninj1,* but were not able to detect any deviations from the reference genome in the *Ninj1* sequence in *Card19^lxcn^* mice. However, *Ninj1* mRNA levels were reduced at baseline in *Card19^lxcn^* BMDMs relative to wild-type BMDMs, and a knockout-validated antibody (41) demonstrated that NINJ1 protein levels were significantly reduced but not entirely absent in *Card19^lxcn^* BMDMs **(Fig. 7E).** To directly test whether loss of NINJ1 in the *Card19^Lxcn^* BMDMs might be responsible for the defect in cell lysis, we reconstituted *Card19^lxcn^* iBMDMs with *Ninj1* and electroporated with LPS. Critically, *Ninj1* reconstitution restored cell lysis to wildtype levels, similarly to the observations for *Ninj1^-/-^* cells **(Fig. 7F, S4B).** Altogether, our findings indicate that in addition to loss of CARD19, *Card19^Lxcn^* mice also have substantially reduced NINJ1 protein levels, most likely as a result of *Card19* targeting, leading to a defect in cell lysis consistent with the loss of NINJ1 function.

### *Card19^lxcn^* mice exhibit increased susceptibility to *Yersinia*

To assess whether the impact on cell death was biologically relevant during an *in vivo* infection in which cell death is important for systemic bacterial control, we infected *Card19^lxcn^* mice with the gram-negative bacterial pathogen *Yersinia pseudotuberculosis* (*Yp*). Consistent with findings that mice lacking essential mediators of cell death are more susceptible to bacterial infection (49, 79–81), *Card19^lxcn^* mice were significantly more susceptible to oral *Yp* infection and had a defect in controlling systemic *Yp* tissue burdens **(Fig. 8A and 8B)**. *Yp-*infected *Card19*^lxcn^ mice exhibited increased splenomegaly **(Fig. 8C and 8D)**, suggesting either failure to control infection and elevated recruitment of inflammatory cells to the spleen, or a failure of innate cells to undergo cell death during bacterial infection in vivo. *Card19^lxcn^* mice had similar levels of serum IL-6 and IL-12, although they had increased levels of TNF **(Fig. 8E)**. Altogether these data indicated that NINJ1-mediated cell lysis is not important for control of cell-intrinsic cytokine production but contributes to host defense against bacterial infection.

**Fig. 8.**
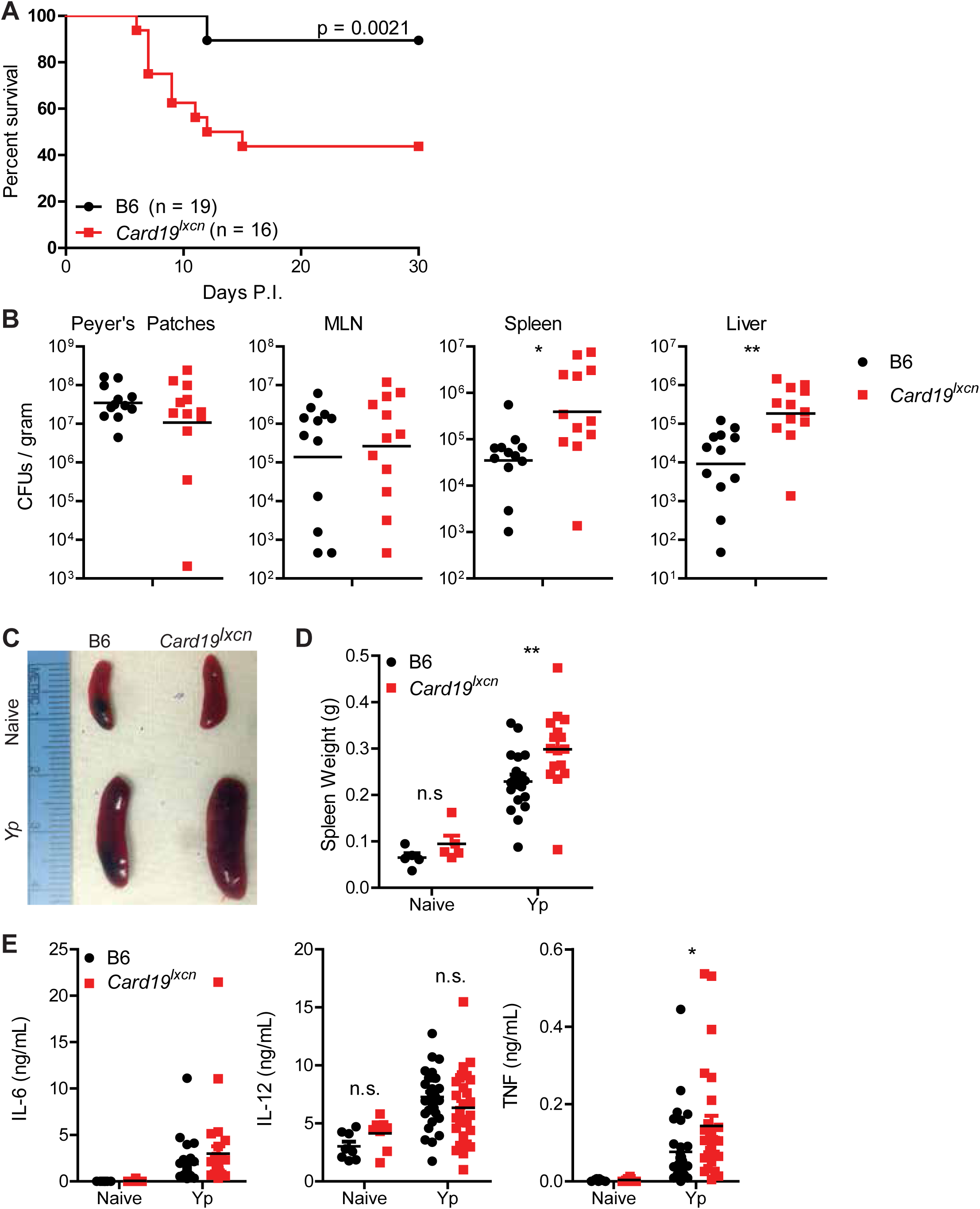
NINJ1 mediates anti-*Yersinia* host defense. (A) Survival of B6 and *Card19*^lxcn^ mice following oral infection 10^8^ CFUs of strain IP2777 *Yp*. Data are pooled from two independent experiments that each gave similar results. Log-rank (Mantel-Cox) Survival test. (B) Bacterial CFUs per gram tissue in Peyer’s Patches, MLN, Spleen and Liver in B6 and *Card19*^lxcn^ mice seven days post-oral infection with *Yp.* Representative of two (PP) or four (MLN, Spleen, Liver) independent experiments. Unpaired Student’s *t*-test. ** p < 0.01, * p < 0.05. (C) Representative images of naive and *Yp*-infected B6 and *Card19*^lxcn^ spleens, seven days post-infection Representative of three independent experiments. (D) Quantification of naive and infected spleen weights. Data are pooled from three independent experiments with similar results. Naive (n=5), *Yp* (B6 n=19, *Card19*^lxcn^ n=16) 2-way ANOVA with Bonferroni multiple comparisons post-test. ** p < 0.01. (E) Serum cytokines from naive and *Yp* infected B6 and *Card19*^lxcn^ mice. Data are pooled from four independent experiments with similar results. Naïve (n=8), *Yp* (B6 n=31, *Card19^lxcn^* n=28) 2-way ANOVA with Bonferroni multiple comparisons post-test. * p < 0.05, n.s. not significant.

## Discussion

Our study has revealed that *Card19^lxcn^* BMDMs are resistant to cell lysis during both pyroptosis and caspase-8-mediated apoptosis while maintaining caspase activation and release of IL-1 family cytokines. This data highlights the existence of a regulatory checkpoint downstream of gasdermin processing that controls the cell fate choice to undergo terminal cell lysis. Until relatively recently, caspase-dependent cleavage of gasdermin D, and the subsequent release of the gasdermin D N-terminal pore-forming domain was viewed as the terminal step in lytic cell death. However, some cell types or stimuli trigger IL-1β release without cell lysis, indicating that these are distinct cellular responses that can be decoupled (28, 29, 31, 82). Moreover, the recent observation that GSDMD-dependent release of IL-1β is regulated by charge-charge interactions highlights the selective nature of the GSDMD pore. That gasdermin-dependent secretion of IL-1 cytokines can be uncoupled from cell lysis further points to regulatory steps subsequent to cleavage and activation of gasdermin D that can be engaged to promote or limit cell lysis.

While our initial observations that *Card19^lxcn^* macrophages were resistant to cell lysis despite having wild-type levels of GSDMD processing suggested that CARD19 might regulate cell lysis, two CRISPR/Cas9-generated lines of mice that we generated directly on a B6 background (*Card19^ΔCARD^* and *Card19^Null^*) did not recapitulate this phenotype. Moreover, expressing CARD19 in *Card19^lxcn^* cells did not complement the lysis phenotype. These findings indicated that a CARD19-independent factor that was presumably affected in *Card19^lxcn^* mice was most likely responsible for the resistance of *Card19^lxcn^* cells to cell lysis. Intriguingly, *Ninj1* which is immediately upstream of *Card19* on chromosome 13, regulates cell lysis in response to pyroptotic and apoptotic stimuli (41). *Ninj1* encodes a protein whose function was previously linked to axonal guidance, and was identified in a forward genetic screen for regulators of plasma membrane rupture (41). The phenotype of *Ninj1*^-/-^ cells is strikingly similar to that of *Card19^lxcn^* BMDMs (41). Although we were unable to identify any mutations in the NINJ1 coding sequence or intronic splice acceptor that would indicate a potential defect in NINJ1 in *Card19^lxcn^* mice, *Card19^lxcn^* BMDMs have substantially diminished levels of NINJ1 gene and protein expression, implicating NINJ1 as the likely driver for the phenotype in the *Card19^lxcn^* line. Furthermore, reconstitution of *Ninj1* in *Card19^lxcn^* iBMDMs restored wildtype levels of cell lysis following stimulation with LPS. Together, these results indicate that genetic targeting of *Card19* resulted in an off-target effect on NINJ1 expression, leading to impaired terminal cell lysis and anti-*Yersinia* defense.

Unintended off-target effects on neighboring genes following genetic targeting have previously been reported. Notably, insertion of a PGK-Neo cassette into the granzyme B locus and β-globin locus control regions was found to disrupt multiple genes in the surrounding area (83). Targeting of exons 13-16 of SH3 and multiple ankyrin repeat domains 3 (Shank3B) with a neomycin resistance cassette resulted in altered expression of neighboring genes as well as increased expression of exons 1-12 and the formation of an unusual Shank3 isoform not expressed in wildtype mice (84). Furthermore, RNA-seq analysis of target and neighboring gene expression in a number of homozygous mutants revealed increased frequency of downregulated neighboring genes that would be expected due to local transcription dysregulation (85). Our finding that genetic targeting of *Card19* resulted in decreased expression of *Ninj1* are consistent with these reports. Our findings underscore the complexity of genetic manipulation in mice and the importance of complementation analyses in validating knockout phenotypes.

In addition to identification of NINJ1 as a regulator of cell lysis, Evavold *et al*. recently uncovered the Ragulator/Rag/mTORC pathway in a genetic screen for factors required in GSDMD-dependent pore formation (86). Intriguingly, the Ragulator complex acts downstream of GSDMD cleavage but upstream of NINJ1-induced cell lysis, as Rag was required for pore formation, whereas NINJ1 acts subsequent to pore formation to induce membrane rupture and ultimate cytolysis (41, 86). Thus, in contrast to initial models proposing that release of the N-terminal portion of GSDMD was necessary and sufficient to trigger cell lysis, these findings collectively support a multi-step regulated process, downstream of GSDMD cleavage, that ultimately triggers lytic rupture of the cell.

How terminal cell lysis occurs downstream of GSDMD cleavage or plasma membrane insertion remains enigmatic. The mitochondrial protein SARM1 was reported to regulate cell lysis in response to NLRP3 inflammasome activation independently of IL-1β cytokine release. Mitochondrial disruption is a feature of both lytic and apoptotic forms of cell death, and the N-terminal GSDMD fragment is recruited to the mitochondrial membrane as well (73, 87). Unexpectedly, in our hands, primary macrophages generated from four independent lines of *Sarm1^-/-^* BMDMs had wild-type levels of cell lysis and IL-1β cytokine release. Moreover, *Sarm1* expression in BMDMs was fairly low in our RNA-seq analysis, and was further reduced upon LPS stimulation. Whether passenger mutations, which have been reported in *Sarm1^-/-^* murine lines (75), may account for the reported phenotype of *Sarm1^-/-^* BMDMs, remains to be determined. Precisely how cell lysis can be uncoupled from IL-1 cytokine release remains a key unanswered question. BMDMs from *Card19^lxcn^*, *Ninj1^-/-^*, and *Rag^-/-^* mice all display reduced cell lysis despite substantial levels of IL-1β cytokine release, similar to the hyperactivation phenotype previously described by Kagan and colleagues (30, 31). However, we observed that BMDMs induced to undergo hyperactivation following infection with Δ*oatA* mutant *S. aureus* exhibit reduced levels of total caspase-1 and GSDMD cleavage, suggesting that the regulated step in hyperactivation occurs at the level of initial inflammasome complex assembly or caspase processing, rather than at the level of GSDMD pore formation.

In addition to demonstrating that terminal cell lysis is a regulated step downstream of GSDMD activation, our study extends the understanding of the regulation of lytic cell death in antibacterial host defense. *Card19^lxcn^* mice showed significantly increased susceptibility to *Yersinia* infection, with higher systemic burdens and reduced ability to survive infection, consistent with our previous studies that the RIPK1-Casp8-dependent cell death pathway promotes cytokine production from uninfected bystander cells. Altogether, these studies highlight cellular rupture as a component of inflammatory responses, independent of IL-1 cytokine release, that contributes to robust antimicrobial immune defense.

## Acknowledgements

We thank Sunny Shin, Jess Doerner, and Daniel Sorobetea for editorial suggestions, and members of the Shin and Brodsky labs, Nobuhiko Kayagaki, and Vishva M. Dixit for scientific discussion. We thank Russell Vance and Isabella Rauch (UC Berkeley) for *Gsdmd^-/-^* bone marrow. We thank Adriano Aguzzi (University Hospital Zurich) for providing the *Sarm1^-/-^* mice (*Sarm1(MSD)*) and Adolfo Garcia-Sastre (Icahn School of Medicine at Mount Sinai) for providing SARM1 knockout bone marrow (*SARM1(AGS)* and *SARM1(AD)*) including wild-type controls. We thank David Sykes (Harvard University) for providing us with ER-HoxB8 reagents.

## Funding

This work was supported by NIH grants AI125924, AI128530, and the Burroughs Wellcome Fund Investigator in the Pathogenesis of Infectious Disease award to I.E.B.

## Competing Interests

Opher S. Kornfeld and Bettina L. Lee are employees of Genentech.

## Data and materials

Anti-caspase-1 antibody and *Ripk3^-/-^* mice were provided by Dr. Vishva M. Dixit (Genentech). *Gsdmd^-/-^* mice were provided by Dr. Russell Vance (University of California, Berkeley). Requests for resources and reagents should be directed to Igor E. Brodsky.

## Materials and Methods

All reagents and resources are listed in Table S5.

### Generation of Card19^ΔCARD^ mice

*Card19^ΔCARD^* mice were produced by the CRISPR/Cas9 Mouse Targeting Core (PSOM-UPenn) using CRISPR/Cas9 technology (flanking sgRNA and Cas9mRA microinjected in 1-cell mouse embryos), and a targeting strategy resulting in removal of exons 2-3. The resulting CARD19 deletion removes amino acids XX including the entire CARD and giving a predicted protein product of 87 AA in length.

CARD19 5’ Target Sequence: GAGATACTGGTGGGACCGAAGGG.

CARD19 3’ Target Sequence: TTGGTCACACTTCGCTGAGATGG.

Mice were backcrossed onto C57BL/6J for at least 2 consecutive generations.

### Generation of *Card19^Null^* mice

*Card19^Null^* mice were produced by the CRISPR/Cas9 Mouse Targeting Core (PSOM-UPenn) using CRISPR/Cas9 technology (flanking sgRNA and Cas9mRA microinjected in 1-cell mouse embryos), and a targeting strategy resulting in deletion of exon 1, resulting in the premature appearance of a stop codon and degradation by nonsense-mediated decay.

CARD19 5’ Target Sequence: TAGGCGAAGGGACGCCGACCCGG.

CARD19 3’ Target Sequence: GTTTCTGGTACGAGCTGGCAGGG.

Mice were backcrossed onto C57BL/6J for at least 2 consecutive generations.

### Differentiation of murine bone marrow-derived macrophages (BMDMs)

Bone marrow derived macrophages were isolated and differentiated as previously described (88). Briefly, isolated bone marrow cells from 6-10 week old male and female mice were grown at 37°C, 5% CO_2_ in 30% macrophage media (30% L929 fibroblast supernatant, complete DMEM). BMDMs were harvested in cold PBS on day 7, and replated in 10% macrophage media onto tissue culture (TC)-treated plates and glass coverslips in TC-treated plates. *Sarm1(AGS3)^-/-^*, *Sarm1(AGS12)^-/-^*, and *Sarm1(AD)^-/-^* were previously described and provided by Adolfo-Garcia Sastre (Icahn School of Medicine at Mount Sinai) along with wildtype controls (74, 75, 89).

### Bacterial culture and infection conditions

Bacteria strains used include *Yersinia pseudotuberculosis* (*Yp*) strain IP2666, *Yp* ΔEJK (lacking Yop effector proteins YopE, YopJ, and YopK), *Salmonella enterica* serovar Typhimurium strain SL1344, *Shigella flexneri* M90T, DH5α *E. coli*, and Δ*oatA Staphylococcus aureus*. Bacterial strains were grown as previously described (26, 49, 71). Briefly, bacteria were grown with aeration and specific antibiotics at 28°C (*Yersinia*, irgasan) or 37°C (*Salmonella*, streptomycin, *E. coli*, none, *S. aureus*, kanamycin). *Shigella* was grown with aeration and without antibiotics at 37°C overnight in Tryptic Soy Broth. *Yersinia* strains were induced prior to infection by diluting the overnight culture 1:40 in 3 mL of inducing media (2xYT broth, 20 mM Sodium Oxalate, 20 mM MgCl_2_). Inducing culture were grown at 28°C for 1 hour and shifted to 37°C for two hours with aeration. *Salmonella* strains were induced prior to infection by diluting the overnight culture 1:40 in 3 mL inducing media (LB broth, 300 mM NaCl), and grown standing for 3 hours at 37°C. Outgrowth cultures of *Shigella* were grown from overnight cultures for 2 hours at 37°C with aeration. Δ*oatA S. aureus* was grown overnight in Todd Hewitt Broth at 37°C with aeration (26). Bacterial growth was measured using OD_600_ on a spectrophotometer. Bacteria were pelleted, washed, and resuspended in DMEM or serum-free media for infection. *In vitro* infections were performed at MOI 10 (*Yp* and *S.* Tm., fractionation and microscopy), MOI 20 (*Yp, ΔEJK* and *S.* Tm., all other assays) or MOI 30 (*E. coli*, LDH) unless otherwise noted. Δ*oatA* infections were performed at MOI 30 and 60. Thioglycolate *in vitro* infections were performed at MOI 10. Gentamicin (100 μg/mL) was added one hour post infection for all infections.

### Mouse strains

*Card19^lxcn^* mice were previously described (33) and maintained as a breeding line in-house. B6 and *Casp1*/*Casp11*^-/-^ mice obtained from Jackson Laboratories and subsequently maintained as a breeding line in-house. All previously published knockout mouse lines that were used to generate BMDMs are indicated in Table S1. *Gsdmd*^-/-^ BMDMs were previously described (90), provided by Russell Vance, and maintained as a breeding line in-house. *Ripk3^-/-^* mice (91) were a gift of Kim Newton and Vishva Dixit (Genentech) and *Ripk3/Casp8^-/-^* mice (92) were a gift of Doug Green (St. Jude Children’s Hospital). *Casp11^-/-^* mice were originally generated by Junying Yuan (93) (Harvard University) and kindly provided by Tiffany Horng (Harvard University). *Sarm1(MSD)^-/-^* mice were a gift from Adriano Aguzzi (University Hospital Zurich). Breeders were routinely genotyped. Mice were maintained in a specific pathogen-free facility by University Laboratory Animal Resources (ULAR) staff in compliance with University of Pennsylvania Institutional Animal Care and Use Committee approved protocols.

### Generation of immortalized *Card19^lxcn^* myeloid progenitors using ER-HOXB8

*Card19^lxcn^* murine myeloid progenitors were immortalized using the ER-HoxB8 system (76). Bone marrow was harvested from female mice and myeloid progenitors were isolated using a Percoll density gradient. Progenitors were plated in 5% SCF-conditioned media supplemented with 10 ng/mL IL-3 and IL-6 for three days. Progenitors were spinfected on fibronectin coated plates with ER-HoxB8 retrovirus in 25 ug/mL polybrene at 1000g for 90 minutes. Progenitors were supplemented with 0.5 uM estrogen and 10 ng/mL GM-CSF to induce macrophage progenitor differentiation. Progenitors were replated with fresh estrogen and GM-CSF for two days. Retrovirally infected cells were selected for with the addition of 1 mg/mL geneticin for 2-3 days. Progenitors were harvested and passaged until immortalization was confirmed by the death of all uninfected control cells. Mature macrophages were generated from immortalized progenitors by plating cells in 30% macrophage media and growing for 5-6 days. Cells were fed with additional macrophage media on day 3.

### Transduction of *Card19^lxcn^* immortalized progenitors

HEK293T cells were transfected using Lipofectamine 2000 with pCL-Eco retroviral packaging vector and MSCV2.2-CARD19 or empty vector (MSCV2.2). Successful transfection was confirmed by imaging under a widefield as MSCV contains IRES-GFP. *Card19^lxcn^* ER-HoxB8 immortalized progenitors were spinfected with filtered viral supernatants from transfected HEK293T cells at 2500 RPM for 90 minutes. Transduced progenitors were transduced a second time the following day as before to increase infection efficiency. Progenitors were allowed to recover and proliferate prior to 2.5 ug/mL puromycin selection. A 95%+ pure population was isolated by flow cytometry sorting on GFP+. Mature macrophages were generated from immortalized, transduced progenitors by plating cells in 30% macrophage media and growing for 5-6 days, with additional media supplementation on day 3.

### NINJ1 reconstitution

Wildtype, *Ninj1^-/-^* and *Card19^lxcn^* iBMDMs were reconstituted with Ninj1 as previously described (41). Briefly, iBMDMs were co-electroporated with NINJ1/the piggyBac vector BH1.11 and the transposase vector pBo using Neon electroporator. 6.25 ug/mL blasticidin was used for selection.

### Plasmids and constructs

All constructs are listed in Table S1. pcDNA3.1/CARD19-FLAG was obtained from GenScript (Refseq NM_026738.2). CARD19 was amplified and inserted into MSCV2.2 using Gibson cloning. All constructs were confirmed by sequencing prior to experimentation.

### Cell death assays – LDH, PI uptake, & Dextran Dye-150 Release

*LDH*. Triplicate wells of BMDMs were seeded in TC-treated 96 well plates. BMDMs were infected with indicated bacterial strains as indicated above. BMDMs were primed with 100 ng/mL LPS for 3 hours followed by 2.5 mM ATP treatment. BMDMs were treated with 10 uM staurosporine or 100 uM etoposide. BMDMs were pretreated with 100 uM zVAD(OMe)-FMK or 100 ug/mL cycloheximide for 1 hour before treatment with 100 ng/mL LPS. For non-canonical inflammasome cell death, BMDMs were primed with 400 ng/mL Pam3CSK4 for 3 hours and then infected with DH5α *E. coli*, MOI 30 for 16-20 hours. For osmoprotected cell death, 5 mM glycine was added at the time of infection. 100 ug/mL gentamicin was added one hour post treatment to all infectious experimental conditions. BMDMs were primed with LPS (100 ng/ml) for 4 hours and stimulated with nigericin (5 μM). BMDMs were primed with Pam3CSK4 (1 μg/ml) for 4 h and were transfected with 2 μg/ml E. coli O111:B4 LPS with Fugene HD. At indicated time points, plates were spun down at 250g and supernatants were harvested. Sups were combined with LDH substrate and buffer according to the manufacturer’s instructions and incubated in the dark for 35 min. Plates were read on a spectrophotometer at 490 nm. Percent cytotoxicity was calculated by normalizing to maximal cell death (1% triton) and no cell death (untreated cells).

*PI uptake.* Propidium Iodide uptake was performed as previously described (26). Briefly, triplicate wells of BMDMs were seeded in TC-treated black-walled 96 well plates. BMDMs were infected or treated as described above in 50 uL HBSS plus 10% FBS. Propidium Iodide (2x, 10 uM) was added in 50 uL HBSS to each well and incubated for 5 minutes in the dark to allow for stabilization of the signal in the maximal cell death wells (1% triton). PI uptake was detected by fluorescence on a BioTek Synergy HT Multi-Detection Microplate Reader (540/25 excitation, 590/35 emission, sensitivity 35, integration time 1 sec) every 2.5 minutes (*S.* Tm., ATP) or every 10 minutes (*Yp*, Sts) for the indicated time points. Gentamicin (100 ug/mL) was added one hour post infection (*Yp*, Sts). Percent cytotoxicity was calculated as described above.

*Dextran Dye-150 (DD-150) Release*. DD-150 release was performed as previously described (41). Briefly, 1.0 × 10^6^ iBMDMs were loaded with DD-150 (50 mg/mL) and stimulated with 0.5 µg LPS via electroporation in R buffer. Prior to plating, iBMDMs were washed with fresh media. Following stimulation, images of iBMDMs were scanned over 20 h with IncuCyte S3 (Essen BioScience) at 10X magnification.

### Confocal microscopy

BMDMs were seeded as described above, treated with staurosporine for indicated time points, fixed, permeabilized, and blocked. BMDMs were stained for HMGB1 (1:200) at 37°C for 1 hr, Alexa Fluor rabbit 488 (1:4000) at RT for 1 hr, and Hoechst and Phalloidin at RT for 30 min. Slides were imaged as described above, with a single z-plane taken per field. Percent nuclear HMGB1 was quantified by identifying total HMGB1 staining per cell and comparing the overlap with nuclear staining (Hoechst). 65-200 cells were analyzed per genotype and condition. Cloud analysis was completed by counting the number of HMGB1 clouds per field of view with 5-8 fields per condition.

### Caspase 8 activity

BMDMs were seeded as described above in a white-walled 96 well plate and treated with *Yp* or Sts for the indicated times. Z-IETD-fmk (500 uM, SM Biochemicals) was added 1 hour prior to infection/treatment as a control to block caspase-8 activity. Caspase-8 activity was detected using Caspase-8 Glo (Promega) according to the manufacturer’s instructions. Luminescence was read on a Gen5 plate reader.

### Western blots

BMDMs were seeded in TC-treated 24 or 12 well plates. Necrostatin (60 uM, Nec-1) was used to inhibit RIPK1 activity for 1 hour prior to *Yp* infection. Following infection or treatment in serum-free media, supernatants were harvested and TCA precipitated. Briefly, supernatants were spun down to remove cell debris and TCA precipitated overnight at 4°C. Sups were spun down and washed with acetone. Remaining TCA was neutralized with Tris, and the pellet was resuspended in 5x sample buffer (125 mM Tris, 10% SDS, 50% glycerol, 0.06% bromophenol blue, 1% β-mercaptoethanol). BMDMs were lysed in lysis buffer (20 mM HEPES, 150 mM NaCl, 10% glycerol, 1% Triton X-100, 1mM EDTA, pH7.5) plus 1x complete protease inhibitor cocktail and 1x sample buffer (25 mM Tris, 2% SDS, 10% glycerol, 0.012% bromophenol blue, 0.2% β-mercaptoethanol). Lysates and supernatants were boiled and centrifuged at full speed for 5 minutes, and sups were run on 4 −12% polyacrylamide gels and transferred to PVDF. Membranes were immunoblotted using the following primary antibodies: β-Actin (1:5000), Bid (1:500), CARD19 (1:500), Caspase 1 (1:360), Caspase 3 (1:1000), Caspase 8 (1:1000), GSDMD (1:1000), HMGB1 (1:1000), GSDME (1:500), PARP (1:1000), NINJ1, Tubulin, and IL-1β. Species specific HRP-conjugated secondary antibodies were used for each antibody (1:5000). Membranes were developed using Pierce ECL Plus and SuperSignal West Femto Maximum Sensitivity Substrate according to the manufacturer’s instructions. Western blot time-courses were performed in parallel with cytotoxicity assays to accurately interpret protein release before and after overt cell death.

### Cytokine release

Triplicate wells of BMDMs were seeded in TC-treated 48 well plates. All conditions except untreated were primed with 100 ng/mL LPS for 3 hrs or unless otherwise indicated. BMDMs were infected with bacterial strains or treated with 2.5 mM ATP. BMDMs were primed with LPS (100 ng/ml) for 4 hours and stimulated with nigericin (5 μM). IL-1β release were measured at the indicated time points. BMDMs were primed with LPS (100 ng/ml) and TNF release was measured after 4 hours. BMDMs were primed with Pam3CSK4 (1 μg/ml) for 4 h and were transfected with 2 μg/ml E. coli O111:B4 LPS with Fugene HD. IL-1β release were measured after 16 h. ELISA supernatants were added to IL-1α, IL-1β, IL-6, IL-12, and TNF-α capture antibody-coated 96-well plates and incubated at 4°C overnight. Plates were incubated with the appropriate biotinylated antibodies in 1% BSA, followed by streptavidin. ELISA was developed with o-phenylenediamine dihydrochloride in citric acid buffer and stopped with 3M sulfuric acid. Plates were read on a spectrophotometer at 490 nm.

### Hyperactivation

Triplicate wells of BMDMs were seeded in TC-treated 48 well plates (ELISA, LDH), and 12 well plates (western). All conditions except untreated were primed with 1 ug/mL LPS for 4 hrs. BMDMs were infected with *S.* Tm. (MOI 20) or *ΔoatA S. aureus* (MOI 30 or 60) as described above. Gentamicin (100 ug/mL) was added 1 hr post infection. Sups and lysates were harvested at 30 min (*S.* Tm.) or 2 and 6 hrs (*ΔoatA*) and analyzed by ELISA for IL-1B release, by LDH for cytotoxicity, or by western blotting for caspase-1, GSDMD, CARD19, and actin (loading control).

### SNP Sequencing

Tails from four *Card19^lxcn^* mice were sent to Dartmouse for a genetic background check. Each sample was interrogated over 5300 SNPs space along the mouse genome with an average density of 0.5 Mbp using an Illumina Infinium Genotyping Assay.

### Whole Exome Sequencing

Genomic DNA was extracted from 1 B6 and two *Card19^lxcn^* BMDMs derived from three independent mice using a DNeasy blood and tissue kit (Qiagen) according to the manufacturer’s instructions. gDNA was shipped to Genewiz for Illumina HiSeq 2×150 bp sequencing. C57BL/6J was used as a reference genome.

### RNA-seq

B6 and *Card19^lxcn^* BMDMs were prepared as above from 3 independent mice per genotype. 1×10^6^ BMDMs were treated with 100 ng/mL for 3 hours or left untreated. RNA was extracted using the RNeasy Plus Mini Kit (Qiagen) according to the manufacturer’s instructions. Sequence-ready libraries were prepared using the Illumina TruSeq Total Transcriptome kit with Ribo-Zero Gold rRNA depletion (Illumina). Quality assessment and quantification of RNA preparations and libraries were carried out using an Agilent 4200 TapeStation and Qubit 3, respectively. Samples were sequenced on an Illumina NextSeq 500 to produce 75–base pair single end reads with a mean sequencing depth of 35 million reads per sample. Raw reads from this study were mapped to the mouse reference transcriptome (Ensembl; Mus musculus GRCm38) using Kallisto version 0.46.2 (94). Raw sequence data are available on the Gene Expression Omnibus (GEO; accession no. GSE168489)

All subsequent analyses were carried out using the statistical computing environment R version 4.0.3 in RStudio version 1.2.5042 and Bioconductor (95). Briefly, transcript quantification data were summarized to genes using the tximport package (96) and normalized using the trimmed mean of M values (TMM) method in edgeR (97). Genes with <1 CPM in 2 samples were filtered out. Normalized filtered data were variance-stabilized using the voom function in limma (98), and differentially expressed genes were identified with linear modeling using limma (FDR ≤ 0.10; absolute log_2_FC ≥ 1) after correcting for multiple testing using Benjamini-Hochberg. Volcano plots were made using the Enhanced Volcano package (99).

### Mouse infections

Age and sex-matched B6 and *Card19^lxcn^* mice between 8-12 weeks were orally infected with 10^8^ CFU of *Yersinia pseudotuberculosis* strain IP2777 after fasting overnight. Mice were monitored for survival for 30 days or were euthanized using CO_2_ followed by cervical dislocation, in compliance with IACUC approved euthanasia protocols. Blood was harvested by cardiac puncture following death. Serum cytokine levels (IL-6, IL-12, TNF-α) were measured by ELISAs as described above. Spleen, liver, mesenteric lymph nodes, and Peyer’s patches were removed, weighed, homogenized, diluted in PBS, and plated on LB agar plates with irgasan for CFUs.

### Thioglycolate Injections

Age and sex-matched B6 and *Card19^lxcn^* mice between 9-11 weeks were injected intraperitoneally with 4% aged thioglycolate or PBS. 48 hours after injection, mice were euthanized using CO_2_. The peritoneum was lavaged with 10 mL cold PBS and the peritoneal exudate cells were spun down. RBCs were lysed using RBC lysis buffer. PECS were counted using trypan blue exclusion on a hemocytometer and plated in complete DMEM without antibiotics in 96 well plates (1×10^5^ cells for PBS PECS, 0.7×10^5^ cells for TG PECS). Cells were washed with warm PBS to remove non-adherent cells and infected with *Yp*, *S.* Tm. or treated with staurosporine as described above.

### Statistical Analysis

Statistical analysis was completed using GraphPad Prism. Two-tailed Student’s *t* test or paired Student’s *t* test were used for comparisons of two groups. One-way analysis of variance (ANOVA) with pairwise comparisons and Bonferroni post-test correction was used for multiple group comparisons. Repeated-measures ANOVA or paired *t* tests were used for matched samples. Studies were conducted without blinding or randomization. Values of p < 0.05 were considered statistically significant

## Notes

### Competing Interest Statement

The authors have declared no competing interest.

### Summary of Updates

We have provided newly obtained data in Figure 7 and Figure 4 indicating reduced NINJ1 expression in the Card19lxcn mice. This data alters our interpretation of the previous data and has led us to revise the abstract and conclusions of the manuscript.

