## Supporting Information for "Genetic targeting of *Card19* is linked to disrupted *Ninj1 expression*, impaired cell lysis, and increased susceptibility to *Yersinia* infection"

Contents

**Fig. S1. *Card[lxcn]* BMDMs are not deficient for Caspase-1, or Caspase-8 activity**
**Fig. S2. SARM1 does not regulate canonical and non-canonical inflammasome death and cytokine release**
**Fig. S3. The defect in cell death is linked to the *Card19* locus**
**Fig. S4. A six megabase region at the *Card19 locus* remains homozygous for 129SvEvBrd**
**Table S1. BMDM and Murine Sources**
**Table S2. Chromosome 13 SNP Results**
**Table S3. Chromosome 13 Whole Exome Sequencing Results**
**Table S4. RNA-Seq Results: Untreated B6 BMDMs vs. Untreated *Card19[lxcn]* BMDMs**
**Table S5. Key resources and reagents**
References

**Table S1: BMDM and Murine Sources**

| Macrophages | Mice Generation | References |
|---|---|---|
| *Card19*$^{lxcn}$ | 129 ESC and backcrossed to B6 | Rios et al. 2020 |
| *Card19*$^{\Delta CARD}$ | C57BL/6J CRISPR Line | This paper |
| *Card19*$^{null}$ | C57BL/6J CRISPR Line | This paper |
| *Sarm1(MSD)*$^{-/-}$ | 129 ESC and backcrossed to B6 | Szrette et al. 2009 |
| *Sarm1(AD)*$^{-/-}$ | 129 ESC and backcrossed to B6 | Kim et al., 2007, JAX stock #018069 |
| *Sarm1(AGS3)*$^{-/-}$ | C57BL/6J CRISPR Line | Uccellini et al., 2020, JAX stock #034399 |
| *Sarm1(AGS12)*$^{-/-}$ | C57BL/6J CRISPR Line | Uccellini et al., 2020 |

**Table S2: Chromosome 13 SNP Results**

| SNP | Location | B6/129 | Change | Gene ID (ENMUSG) | Gene Name | Function |
|---|---|---|---|---|---|---|
| **rs30100204** | 39337735 | B6 | intergenic variant | | | |
| **rs36514425** | 39958176 | B6 | intergenic variant | | | |
| **rs29228586** | 40345334 | B6 | intron variant | 00000047094 | orofacial cleft 1 candidate | associated with cleft lip |
| **rs50931018** | 42450412 | B6 | intergenic variant | | | |
| **rs29236582** | 42602578 | B6 | intergenic variant | | | |
| **rs29635560** | 42602754 | B6 | intergenic variant | | | |
| **rs13481783** | 42840147 | B6 | intron variant | 00000054728 | phosphatase and actin regulator 1 | motility and cytoskeletal organization |
| **rs46355744** | 43000858 | B6 | intron variant | 00000054728 | | |
| **rs29552103** | 43005082 | B6 | intron variant | 00000054728 | | |
| **rs51682551** | 43032240 | B6 | intron variant | 00000054728 | | |
| **rs3712907** | 43171011 | B6 | intron variant | 00000021368 | Tbc1d7 | cell growth and differentiation |
| **rs29995243** | 43206944 | B6 | intron variant | 00000051335 | glucose-fructose oxidoreductase domain containing 1 | |
| **rs29864465** | 44039283 | B6 | intergenic variant | | | |
| **rs46963560** | 45010065 | B6 | intergenic variant | | | |
| **rs6296954** | 45351934 | B6 | intergenic variant | | | |
| **rs3688207** | 45359288 | B6 | intergenic variant | | | |
| **rs108216631** | 45395035 | B6 | intron variant | 00000038175 | Idol/Mir/Myelip | lipid metabolism |
| **rs29225851** | 46060929 | B6 | intron variant | 00000221550 | predicted gene | |
| **rs3663819** | 46297108 | B6 | intron variant | 00000063529 | stathmin domain containing 1 | cell differentiation |
| **rs6411274** | 47129920 | B6 | intron variant | 00000038068 | ring finger protein 144B , lbrdc2 | E3 ubiquitin ligase |
| **rs29225085** | 47229758 | B6 | intron variant | 00000038068 | ring finger protein 144B , lbrdc2 | E3 ubiquitin ligase |
| **rs29914889** | 47764476 | B6 | intergenic variant | | | |

| | | | | | | |
|---|---|---|---|---|---|---|
| **rs6244558** | 47815991 | B6 | intergenic variant | | | |
| **rs220959940** | 48010713 | B6 | intron variant | 00000047324 | RIKEN cDNA 4931429P17 | |
| **rs29568118** | 48119885 | B6 | intron variant | 00000097622 | RIKEN cDNA A330033J07 | |
| **rs30191571** | 48607862 | B6 | intron variant | 00000038042 | protein tyrosine phosphatase domain containing 1 | |
| | **49340961** | | | **00000037966** | **Ninjurin 1 (NINJ1)** | Terminal Pore Regulation |
| | **49356426** | | | **1110007C09Rik** | **CARD19** | |
| **rs37780795** | 49403959 | B6 | intergenic variant | | | |
| **rs6330796** | 51046190 | B6 | intergenic variant | | | |
| **rs226625541** | 51769402 | 129 | intron variant | 00000021451 | Sema4d | Signaling |
| **rs46751182** | 52014198 | B6 | | 00000793968 | | |
| **rs30070966** | 52018425 | 129 | downstream gene variant | 00000102173 | predicted gene | |
| **rs29231157** | 52019172 | 129 | downstream gene variant | 00000102173 | predicted gene | |
| **rs36690691** | 52951172 | 129 | intergenic variant | | | |
| **rs29529592** | 53233552 | 129 | intron variant | 00000021464 | ntrk2 | |
| **rs49400466** | 53412842 | 129 | intron variant | 00000107008 | predicted gene | |
| **rs30058409** | 53813956 | 129 | intergenic variant | | | |
| **rs29551959** | 54964992 | 129 | intron variant | 00000025876 | unc5h1 | |
| **rs30004717** | 54967613 | 129 | intron variant | 00000025876 | unc5h1 | |
| **rs29927068** | 55063173 | 129 | intron variant | 00000025878 | ubiquitin interaction motif containing 1 | |
| **rs29239941** | 56024612 | 129 | intron variant | 00000114493 | predicted gene | |
| **rs29234727** | 56057642 | 129 | intron variant | 00000114493 | predicted gene | |
| **rs29227915** | 56074012 | 129 | 3' UTR variant | 00000015937 | macroH2A.1 histone | |
| **rs46196633** | 56077106 | 129 | intron variant | 00000015937 | macroH2A.1 histone | |
| **rs3720782** | 56229525 | 129 | intron variant | 00000097361 | RIKEN cDNA 4930550C17 gene | |
| **rs3700819** | 57253190 | B6 | intergenic variant | | | |
| **rs30059311** | 57271612 | B6 | intergenic variant | | | |
| **rs29249300** | 57408045 | B6 | intergenic variant | | | |
| **rs30078817** | 57485175 | B6 | intron variant | 00000056222 | Ticn1/Spock1 | Cell-cell interactions |
| **rs257680153** | 58007116 | B6 | intron variant | 00000014164 | klhl3 | Nephron ion transport |
| **rs13481832** | 58837796 | B6 | splice region variant | 00000055254 | neurotrophic tyrosine kinase, receptor, type 2 | Neuronal homeostasis and development |

| rs30250735 | 59737415 | B6 | downstream gene variant | 00000181528 | RIKEN cDNA 4930528D03 |
|---|---|---|---|---|---|

SNPs from DartMouse SNP genetic background check 10 megabases upstream and downstream of the *Ninj1/Card19* chromosome 13 locus. SNP identification and chromosomal position are indicated. SNPs are indicated as wildtype (B6) or 129SvEvBrd (129). The change, gene ID, gene name, and known function are listed. *Ninj1* and *Card19* are listed for reference.

**Table S3: Chromosome 13 Whole Exome Sequencing Results**

| Gene ID | Gene Name | Position | Change | Function and notes |
|---|---|---|---|---|
| *Spata31* | spermatogenesis associated 31 | 13, 34.21 | R645 to stop | spermatogenesis; testis specific |
| *Nlrp4f* | NLR family, pyrin domain containing 4F | 13, 34.45 | E604Q | organelle development |
| *Gm10324* | predicted gene 10324 | 13, 34.51 | S310C, Y316H, A363V, K458R, F551Y | testis, placenta specific |
| *2410141K09Rik* | RIKEN cDNA 2410141K09 gene | 13, 34.52 | several intergenic region Indels | testis specific |
| *2410141K09Rik* | RIKEN cDNA 2410141K09 gene | 13, 34.52 | intron variant, 153A>G | testis specific |
| *Adcy2* | adenylate cyclase 2 | 13, 35.55 | intergenic region, 6802150A>G | cAMP signaling |
| *Cmya5* | cardiomyopathy associated 5 | 13, 47.81 | A3414P | anchoring protein for PKA, skeletal muscle regeneration |
| **Card*19*** | | **13, 49.35-36** | | |
| *Naip1* | NLR family, apoptosis inhibitory protein 1 | 13, 53.18 | W288L | inhibits apoptosis |
| *Tmem267* | transmembrane protein 267 | 13, 67.25 | intron variant, 1131C>T | testis specific |

**Table S4: RNA-Seq Results: Untreated B6 BMDMs vs. Untreated *Card19*$^{lxcn}$ BMDMs**

| Gene ID | B6 Average | *Card19*$^{lxcn}$ Average | Log Fold Change | Function |
|---|---|---|---|---|
| *Sirt5* | 0.44 | -6.27 | -6.71 | NAD+ regulation |
| *Ninj1* | 6.45 | 4.92 | -1.53 | Plasma membrane rupture regulation |
| *Axl* | 6.59 | 5.51 | -1.07 | Inhibits TLR signaling |
| *Wdfy1* | 3.82 | 4.83 | 1 | Positively regulates TLR signaling |
| *Cxcl14* | 5.47 | 7.14 | 1.67 | Chemokine for innate immune cells |

**Table S5. Key resources and reagents.**

| REAGENT or RESOURCE | SOURCE | IDENTIFIER |
|---|---|---|
| **Antibodies** | | |
| Anti-CARD19 polyclonal antibody | Atlas Antibodies | Cat# HPA010990, RRID:AB_2668400 |
| Caspase-1 antibody | Genetech | N/A |
| Rabbit polyclonal to Caspase-3 | Cell Signaling Technologies | Cat# 9662S, RRID:AB_10694681 |
| Rat Anti-Mouse Caspase-8 Monoclonal Antibody, Clone 1G12 | Enzo Life Technologies | Cat# ALX-804-447-C100, RRID:AB_2050952 |
| Anti-DFNA5/GSDME antibody [EPR19859] – N-terminal | Abcam | Cat# ab215191, RRID:AB_2737000 |
| Rabbit monoclonal [EPR19828] to GSDMD | Abcam | Cat# ab209845 |

| | | |
|---|---|---|
| Rabbit Polyclonal to HDAC1 | Cell Signaling Technologies | Cat# 2062, RRID:AB_2118523 |
| HA-Tag (C29F4) Rabbit mAb | Cell Signaling | Cat#3724T |
| HMGB1 antibody - ChIP Grade | Abcam | Cat# ab18256, RRID:AB_444360 |
| Purified Mouse anti-Ninjurin GNE 425 | Genetech | N/A |
| Monoclonal Anti-alpha-Tubulin antibody produced in mouse | Sigma-Aldrich | Cat# T5168, RRID:AB_477579 |
| Goat Anti-Rabbit IgG (H+L) Antibody, Alexa Fluor 488 Conjugated | Molecular Probes | A-11008, RRID:AB_143165 |
| Goat anti-Mouse IgG (H+L) Cross-Adsorbed Secondary Antibody, Alexa Fluor 514 | Thermo Fisher Scientific | Cat# A-31555, RRID:AB_2536171 |
| Alexa Fluor® 647 Phalloidin antibody | Thermo Fisher Scientific | Cat# A22287, RRID:AB_2620155 |
| Mouse Anti-beta-Actin Monoclonal Antibody, Unconjugated, Clone AC-74 | Sigma-Aldrich | Cat# A2228, RRID:AB_476697 |
| Peroxidase-AffiniPure Goat Anti-Rat IgG (H+L) (min X Hu,Bov,Hrs,Rb Sr Prot) antibody | Jackson Labs | Cat# 112-035-143, RRID:AB_2338138 |
| Peroxidase-AffiniPure Goat Anti-Rabbit IgG (H+L) (min X Hu,Ms,Rat Sr Prot) antibody | Jackson Labs | Cat# 111-035-144, RRID:AB_2307391 |
| Anti-mouse IgG, HRP-linked Antibody | Cell Signaling Technologies | Cat# 7076, RRID:AB_330924 |
| IL-1a antibody | BD Biosciences | Cat# 550604, RRID:AB_393776 |
| Biotin anti-mouse IL-1a antibody | BioLegend | Cat# 512504, RRID:AB_2124220 |
| Anti-Mouse/Rat IL-1 beta Purified 500 ug antibody | Thermo Fisher Scientific | Cat# 14-7012-85, RRID:AB_468397 |
| Anti-Mouse IL-1 beta Biotin 500 ug antibody | Thermo Fisher Scientific | Cat# 13-7112-85, RRID:AB_466925 |
| Rat Anti-IL-6 Monoclonal Antibody, Unconjugated, Clone MP5-20F3 | BD Biosciences | Cat# 554400, RRID:AB_398549 |
| Rat Anti-IL-6 Monoclonal Antibody, Biotin Conjugated, Clone MP5-32C11 | BD Biosciences | Cat# 554402, RRID:AB_395368 |
| Rat Anti-IL-12 (p40 / p70) Monoclonal Antibody, Unconjugated, Clone C15.6 | BD Biosciences | Cat# 551219, RRID:AB_394097 |
| Rat Anti-IL-12 (p40 / p70) Monoclonal Antibody, Biotin Conjugated, Clone C17.8 | BD Biosciences | Cat# 554476, RRID:AB_395419 |
| IFN gamma Monoclonal Antibody (AN-18), eBioscience(TM) | Thermo Fisher Scientific | Cat# 14-7313-85, RRID:AB_468472 |
| IFN gamma Monoclonal Antibody (R4-6A2), Biotin, eBioscience(TM) | Thermo Fisher Scientific | Cat# 13-7312-85, RRID:AB_466939 |
| Goat anti-Mouse IgG1 Secondary Antibody, Alexa Fluor 488 conjugate | Thermo Fisher Scientific | Cat# A-21121, RRID:AB_2535764 |
| Goat anti-Mouse IgG1 Cross-Adsorbed Secondary Antibody, Alexa Fluor 647 | Thermo Fisher Scientific | Cat# A-21240, RRID:AB_2535809 |
| Goat anti-Mouse IgG2a Cross-Adsorbed Secondary Antibody, Alexa Fluor 488 | Thermo Fisher Scientific | Cat# A-21131, RRID:AB_2535771 |
| Goat anti-Rabbit IgG (H+L) Highly Cross-Adsorbed Secondary Antibody, Alexa Fluor 647 | Thermo Fisher Scientific | Cat# A-21245, RRID:AB_2535813 |
| Goat anti-Rabbit IgG (H+L) Secondary Antibody, Alexa Fluor 546 | Thermo Fisher Scientific | Cat# A-11010, RRID: AB_143156 |
| Goat anti-Mouse IgG (H+L) Secondary Antibody, Alexa Fluor 546 | Thermo Fisher Scientific | Cat# A-11003, RRID: AB_2534071 |
| **Bacterial and Virus Strains** | | |
| *Yersinia pseudotuberculosis* IP2777 | Stanley Falkow | (1) |
| *Yersinia pseudotuberculosis* IP2666 | Jim Bliska | (2) |
| *Yersinia pseudotuberculosis* IP2666 ΔyopEJK | Erin Zwack, Igor Brodksy | (3) |
| *Salmonella* Typhimurium SL1344 | (4) | |
| *ΔoatA Staphylococcus aureus* | Jonathan Kagan | (5) |

| | | |
|---|---|---|
| *E. coli* DH5α | N/A | |
| ER-HoxB8 Virus | David Sykes | (6) |
| **Biological Samples** | | |
| *Gsdmd*$^{-/-}$ Bone Marrow | Russell Vance | (7) |
| *Mavs*$^{-/-}$ Bone Marrow | Carolina Lopez, Jackson Labs | Cat# 008634 |
| *Casp11*$^{-/-}$ Bone Marrow | Junying Yuan | (8) |
| *Sarm1(AGS3)*$^{-/-}$, *Sarm1(AGS12)*$^{-/-}$, *Sarm1(AD)*$^{-/-}$ Bone Marrow | Adolfo Garcia-Sastre | (9) |
| *Ninj1*$^{-/-}$ *iBMDMs* | Vishva Dixit | (10) |
| **Chemicals, Peptides, and Recombinant Proteins** | | |
| Hoechst | Thermo Fisher Scientific | Cat# 62249 |
| Fluormount G | Thermo Fisher Scientific | Cat# 00-4958-02 |
| Lipopolysaccharide from E Coli | Sigma-Aldrich | Cat# L2880 |
| Pam3CSK4 | Invivogen | Cat# tlrl-pms |
| Cycloheximide from microbial | Sigma-Aldrich | Cat# C7698 |
| zVAD(Ome)-FMK | SM Biochemicals | Cat# SMFMK001 |
| z-IETD-FMK | SM Biochemicals | Cat# SMFMK004 |
| Adenosine 5'-Triphosphate, Disodium Salt | Millipore | Cat# 1191 |
| Gentamicin Sulfate | Sigma-Aldrich | Cat# G1914 |
| Irgasan | Sigma-Aldrich | Cat# 72779 |
| Streptamycin Sulfate | Gold Technologies | Cat# G-400-1 |
| Staurosporine from *Streptomyces sp.* | Sigma-Aldrich | Cat# S5921 |
| Necrostatin (Nec-1) | EMD Chemicals | Cat# 480065 |
| MitoTracker CMXRos | Life Technologies | Cat# M7512 |
| Propidium iodide | Thermo Fisher Scientific | Cat# P3566 |
| ECL Western Blotting Substrate | Thermo Fisher Scientific | Cat# 32106 |
| SuperSignal West Femto Maximum Sensitivity Substrate | Thermo Fisher Scientific | Cat# 34095 |
| Complete, Mini, EDTA-Free Protease Inhibitor Cocktail | Roche | Cat# 11836170001 |
| Streptavidin HRP | BD Biosciences | Cat# 554066 |
| Recombinant Mouse TNF-alpha | BioLegend | Cat# 575206 |
| Recombinant Mouse IFN-gamma protein | eBioscience | Cat# 485-MI-100 |
| Mouse IL-1beta Recombinant Protein | R&D | Cat# 14-8012-80 |
| Recombinant IL-6 Standard | R&D | Cat# 406-ML-005 |
| Recombinant Mouse IL-12 protein | R&D | Cat# 419-ML-010 |
| Recombinant Mouse IL-1 alpha | R&D | Cat# 400-ML-005 |
| Lipofectamine 2000 | Invitrogen | Cat# 11668027 |
| 16% Paraformaldehyde Aqueous Solution, EM Grad | Electron Microscopy Services | Cat# 15710 |
| Mounting Media, PPD in 90% Glycerol | (11) | |
| DAPI | Molecular Probes | Cat# D-1306 |
| Todd Hewitt Broth for microbiology | Sigma-Aldrich | Cat# T1438 |
| Dextran Dye-150 | Millipore Sigma | Cat# FD150S |
| **Commercial Assays** | | |
| Plasma Membrane Protein Extraction Kit | Abcam | Cat# ab65400 |
| LDH Cytotoxicity Detection Kit | Takara Bio | Cat# MK401 |
| CellTiter Glo Luminescent Cell Viability Assay | Promega | Cat# G7571 |
| Plasmid Maxiprep Kit | Qiagen | Cat# 12162 |
| Caspase-Glo 8 Assay | Fisher Scientific | Cat# PRG8201 |
| **Experimental Models: Cell Lines** | | |
| HEK 293T Cells | ATCC | |
| **Experimental Models: Organisms/Strains** | | |
| *Card19*$^{lxcn}$ Mice | Brian Schaefer | (12) |

| | | |
|---|---|---|
| *Card19*$^{\Delta CARD}$ Mice | This paper | |
| *Card19*$^{Null}$ Mice | This paper | |
| C57BL/6J Mice (B6) | Jackson Labs | Cat# 000664 |
| *Casp1/Casp11*$^{-/-}$ Mice | Jackson Labs | Cat# 016621 |
| *Ripk3/Casp8*$^{-/-}$ Mice | Doug Green | (13) |
| *Sarm1(MSD)*$^{-/-}$ | Adriano Aguzzi | (14) |
| *Ripk3*$^{-/-}$ Mice | Kim Newton, Vishva Dixit | (15) |
| **Oligonucleotides** | | |
| *Card19*$^{lxcn}$ Wt genotyping primers, (297 bp)<br>CATGGATGTACAGAGCTCGGTAA,<br>CGTTGCCCTGGAGACACAGTATT | IDT | This paper |
| *Card19*$^{lxcn}$ Knockout genotyping primers (281 bp)<br>CGGAATTGATCCCGCTCGAA,<br>CGTTGCCCTGGAGACACAGTATT | IDT | This paper |
| *Card19*$^{\Delta CARD}$ Sequencing Primer (250 bp)<br>CTTGGGAAAAGTGTGGCTTTTGT,<br>TCCTCCAGTCTGTCCATGTGGGGATTTT | Sigma | This paper |
| *Card19*$^{Null}$ Sequencing primer (298 bp)<br>TCGGTTTCTTCATCCAGGAG,<br>GAGGCAGCCACTGGGTATAA | Sigma | This paper |
| **Recombinant DNA** | | |
| pcDNA3.1+/CARD19-FLAG | GenScript | Cat# OMu021914D |
| pcDNA3.1+ | Igor Brodsky | |
| pMSCV2.2 | Igor Brodsky | |
| pMSCV2.2/CARD19 | This paper | |
| pCL-Eco | Igor Brodsky | |
| mNINJ1/BH1.11 | Vishva Dixit | (10) |
| BH1.11 | Vishva Dixit | (10) |
| pBO | Vishva Dixit | (10) |
| **Software and Algorithms** | | |
| FIJI | (16) | https://imagej.net/Fiji/Downloads |
| Volocity 6.3 | PerkinElmer | http://cellularimaging.perkinelmer.com/downloads/detail.php?id=14 |
| Prism 5.0 | GraphPad | https://www.graphpad.com/scientific-software/prism/ |
| R version 4.0.3 | R | https://www.r-project.org/ |
| RStudio version 1.2.5042 | RStudio | https://rstudio.com/ |
| Other | | |

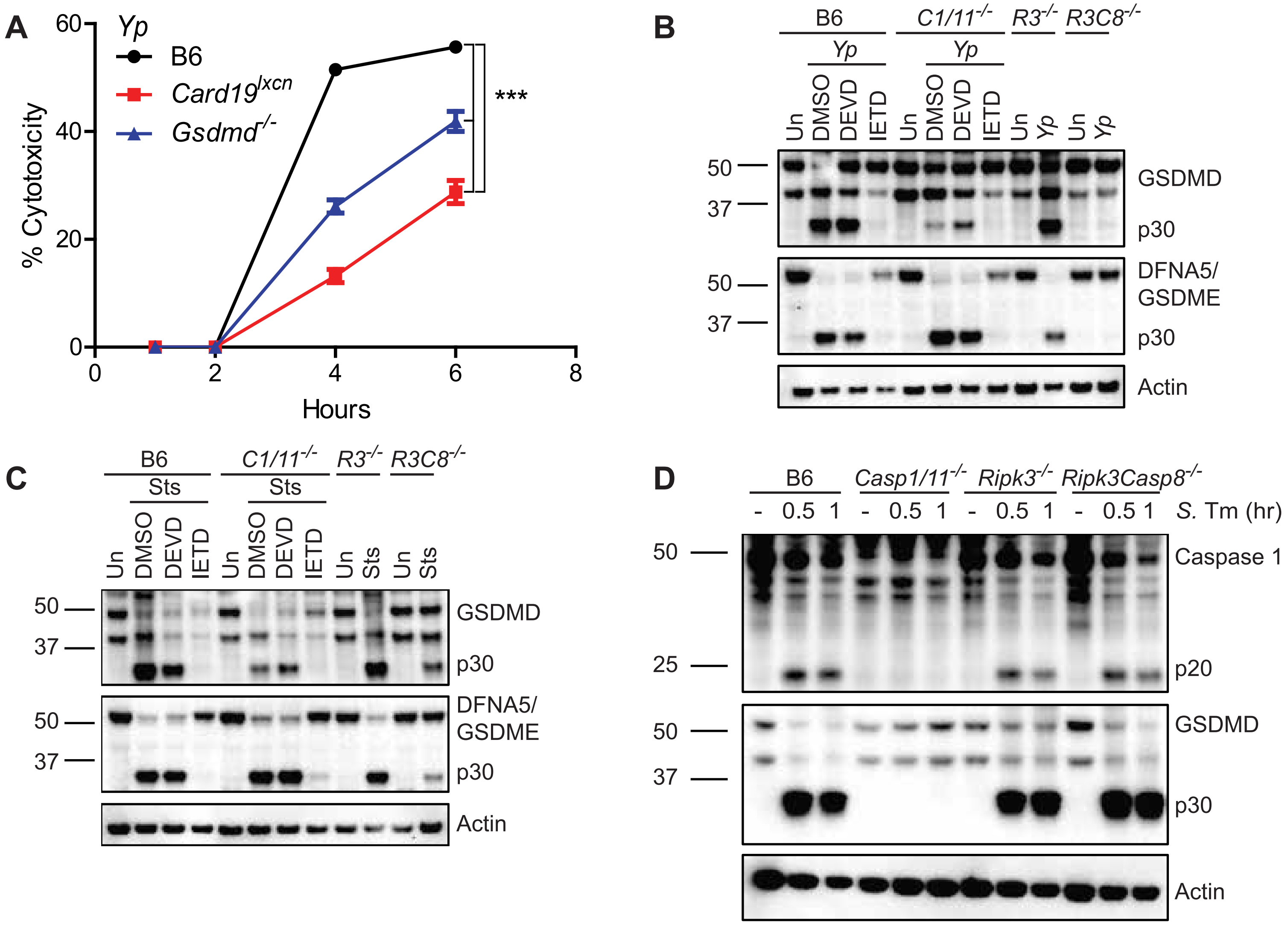

A
Yp
B6
Card19lxcn
Gsdmd-/-
% Cytotoxicity
Hours
***
B
B6
C1/11-/-
R3-/-
R3C8-/-
Yp
Un
DMSO
DEVD
IETD
GSDMD
p30
DFNA5/
GSDME
Actin
C
Sts
D
Casp1/11-/-
Ripk3-/-
Ripk3Casp8-/-
S. Tm (hr)
Caspase 1
p20

S1

**Fig. S1. Card$^{lxcn}$ BMDMs are not deficient for Caspase-1, or Caspase-8 activity**

(A) B6, *Card19*$^{lxcn}$, and *Gsdmd*$^{-/-}$ BMDMS were infected with *Yp*. Cell death was assayed by LDH release. Representative of three independent experiments.

(B-D) B6, Casp1/11$^{-/-}$, *Ripk3*$^{-/-}$, or *Ripk3*$^{-/-}$/*Casp8*$^{-/-}$ BMDMs were left uninfected (Un) (B) infected with *Yp*, (C) treated with sts or (D) infected with *S.* Tm in the presence of DMSO, the caspase-3/7 inhibitor DEVD, and the caspase-8 inhibitor IETD. Lysates were harvested (B, C) 3 hours or (D) 0.5 and 1 hour post treatment and analyzed by western blotting for GSDMD, DFNA5/GSDME, Caspase-1, and actin (loading control). Blots representative of two or three independent experiments.

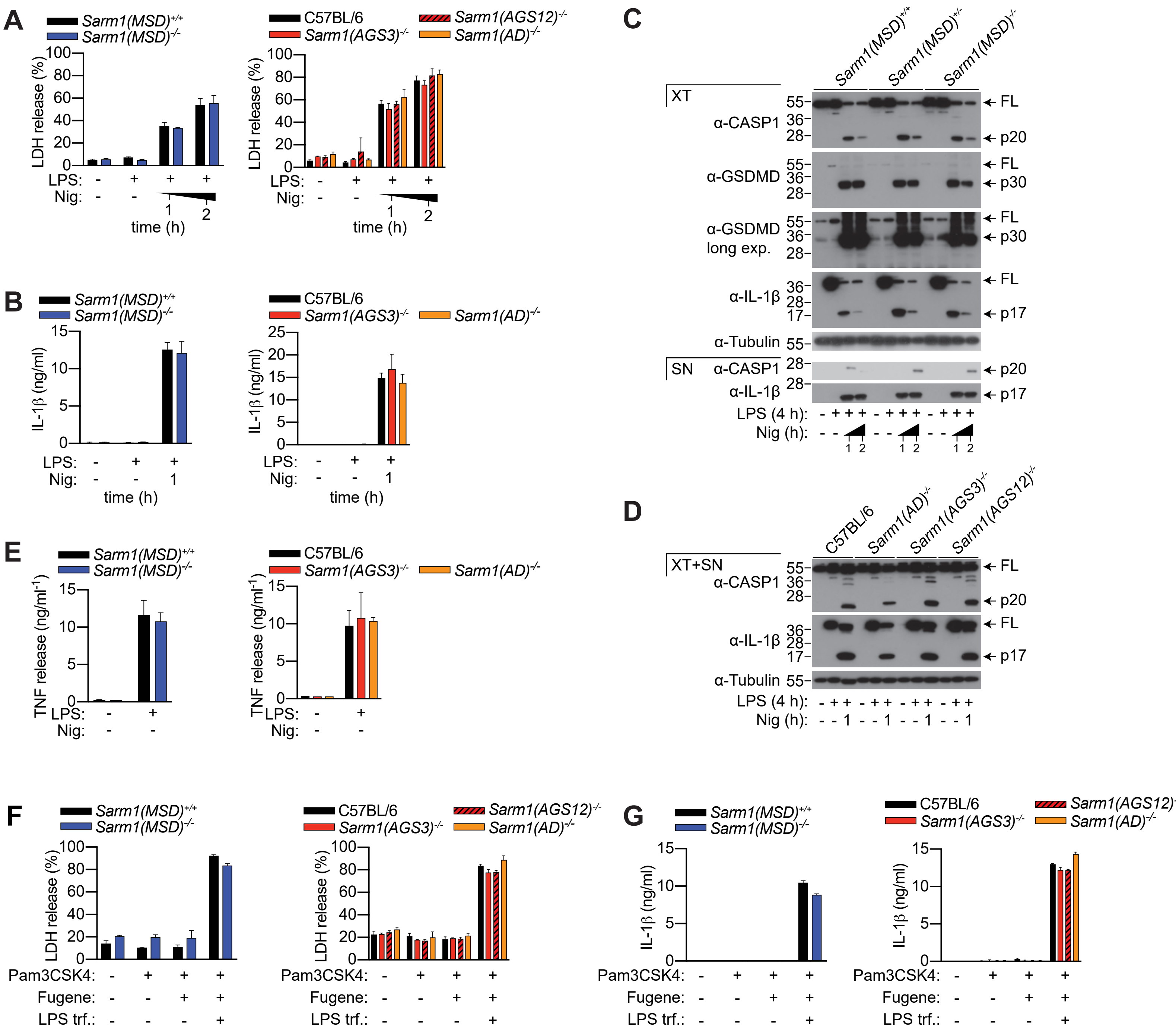
A
Sarm1(MSD)+/+
Sarm1(MSD)-/-
LDH release (%)
LPS:
Nig:
time (h)
C57BL/6
Sarm1(AGS3)-/-
Sarm1(AGS12)-/-
Sarm1(AD)-/-
B
IL-1β (ng/ml)
C
Sarm1(MSD)+/+
Sarm1(MSD)+/-
Sarm1(MSD)-/-
XT
α-CASP1
α-GSDMD
α-GSDMD long exp.
α-IL-1β
α-Tubulin
SN
FL
p20
p30
p17
LPS (4 h):
Nig (h):
D
XT+SN
E
TNF release (ng/ml-1)
F
Pam3CSK4:
Fugene:
LPS trf.:
G

S2

**Fig. S2. SARM1 does not regulate canonical and non-canonical inflammasome death and cytokine release**

(A to D) *Sarm1(MSD)*$^{-/-}$, *Sarm1(MSD)*$^{+/-}$, *Sarm1(MSD)*$^{+/+}$, C57BL/6, *Sarm1(AGS3)*$^{-/-}$, *Sarm1(AGS12)*$^{-/-}$ and *Sarm1(AD)*$^{-/-}$ BMDMs were primed with LPS (100 ng/ml) for 4 hours and stimulated with nigericin (5 μM). (A) LDH and (B) IL-1β release were measured at the indicated time points. (C) Supernatant and cell extract or (D) mixed supernatant and cell extract were examined by immunoblotting at the indicated time points.

(E) BMDMs were primed with LPS (100 ng/ml) and TNF release was measured after 4 hours.

(F and G) BMDMs were primed with Pam3CSK4 (1 μg/ml) for 4 h and were transfected with 2 μg/ml E. coli O111:B4 LPS with Fugene HD. (F) LDH and (G) IL-1β release were measured after 16 h.

(A and B, E-G) Data are mean + SD of triplicate cell stimulation and is representative of two to four independent experiments. Immunoblots are representative of two independent experiments.

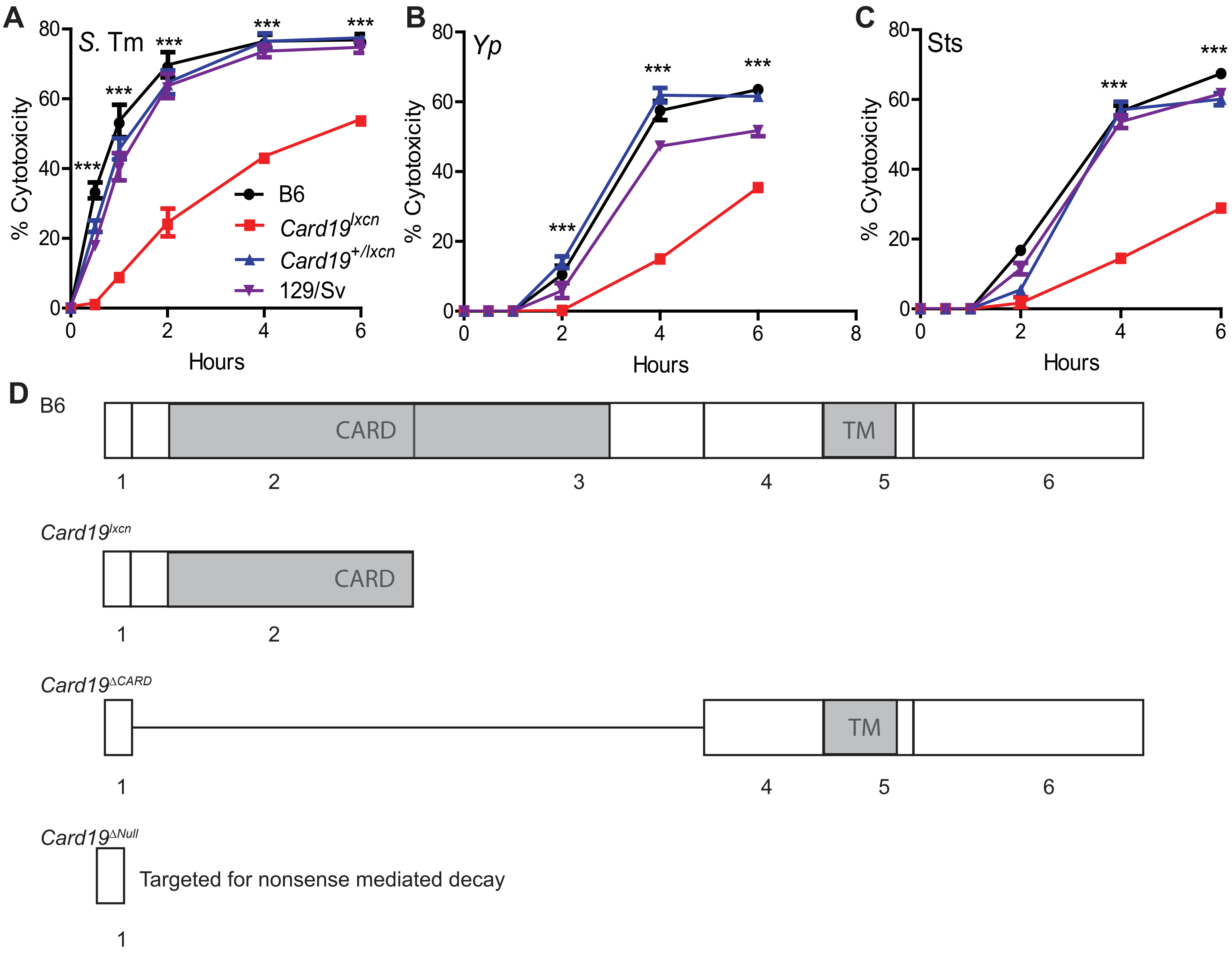
A
S. Tm
B
Yp
C
Sts
% Cytotoxicity
Hours
***
B6
Card19lxcn
Card19+/lxcn
129/Sv
D
B6
CARD
TM
1
2
3
4
5
6
Card19lxcn
Card19ΔCARD
Card19ΔNull
Targeted for nonsense mediated decay

S3

**Fig. S3: The defect in cell death is linked to the *Card19* locus**

(A-C) B6, Card19$^{lxcn}$, Card19$^{+/lxcn}$, and 129/Sv BMDMs were treated with (A) *S.*Tm, (B) *Yp* or (C) sts. Cell death was assayed by LDH release at indicated time points. Representative of three independent experiments.

(D) CARD19 exon schematic showing each independent CARD19 mouse line with the respective CARD19 product.

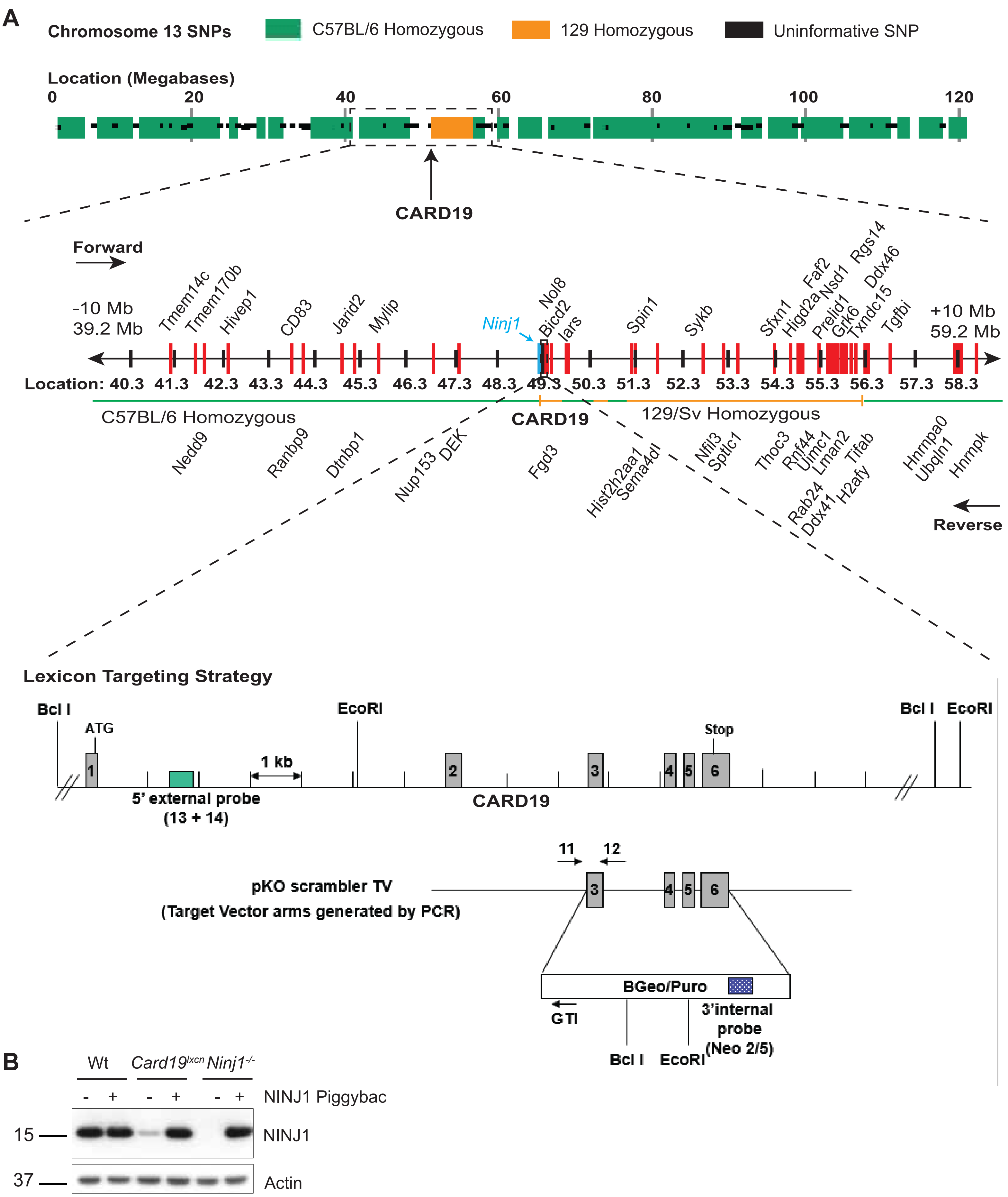
A
Chromosome 13 SNPs
C57BL/6 Homozygous
129 Homozygous
Uninformative SNP
Location (Megabases)
0
20
40
60
80
100
120
CARD19
Forward
-10 Mb
39.2 Mb
Tmem14c
Tmem170b
Hivep1
CD83
Jarid2
Mylip
Ninj1
Nol8
Bicd2
Iars
Spin1
Sykb
Sfxn1
Higd2a
Faf2
Nsd1
Rgs14
Prelid1
Grk6
Txndc15
Ddx46
Tgfbi
+10 Mb
59.2 Mb
Location: 40.3 41.3 42.3 43.3 44.3 45.3 46.3 47.3 48.3 49.3 50.3 51.3 52.3 53.3 54.3 55.3 56.3 57.3 58.3
C57BL/6 Homozygous
CARD19
129/Sv Homozygous
Nedd9
Ranbp9
Dtnbp1
Nup153
DEK
Fgd3
Hist2h2aa1
Sema4dl
Nfil3
Sptlc1
Thoc3
Rnf44
Uimc1
Lman2
Tifab
Rab24
Ddx41
H2afy
Hnrnpa0
Ubqln1
Hnrnpk
Reverse
Lexicon Targeting Strategy
Bcl I
ATG
EcoRI
Bcl I
EcoRI
Stop
1
2
3
4
5
6
1 kb
5' external probe
(13 + 14)
CARD19
11
12
pKO scrambler TV
(Target Vector arms generated by PCR)
3
4
5
6
BGeo/Puro
GTI
3'internal
probe
(Neo 2/5)
Bcl I
EcoRI
B
Wt
Card19lxcn
Ninj1-/-
- + - + - +
NINJ1 Piggybac
15
NINJ1
37
Actin

S4

**Fig. S4: A six megabase region at the *Card19 locus* remains homozygous for 129SvEvBrd**

(A) Chromosome 13 with tested SNPs from DartMouse genetic background check. C57BL/6 SNPs are in green, 129SvEvBrd SNPs are in yellow, and uninformative SNPs (i.e. not all samples gave identical results) are in black. The 10 megabase region on either side of the *Card19* locus is zoomed in below it with chromosomal locations noted in bold numbers and black notches. Genes in red are expressed in macrophages. *Ninj1* is in blue. The six megabase region highlighted by the yellow bar is homozygous for 129SvEvBrd. The green regions are homozygous for C57BL/6. A zoomed in region at *Card19* is highlighted with the original Lexicon Targeting strategy.

(B) Wildtype, *Card19*$^{lxcn}$*, and Ninj1*$^{-/-}$ iBMDMs were reconstituted with NINJ1/BH1.11 piggyBac or empty vector. Lysates were harvested and run on SDS-PAGE gel and probed for NINJ1 and actin (loading control).
